# Pediatric human nose organoids demonstrate greater susceptibility, epithelial responses, and cytotoxicity than adults during RSV infection

**DOI:** 10.1101/2024.02.01.578466

**Authors:** Gina M. Aloisio, Divya Nagaraj, Ashley M. Murray, Emily M. Schultz, Trevor McBride, Letisha Aideyan, Erin G. Nicholson, David Henke, Laura Ferlic-Stark, Anubama Rajan, Amal Kambal, Hannah L. Johnson, Elina Mosa, Fabio Stossi, Sarah E. Blutt, Pedro A. Piedra, Vasanthi Avadhanula

## Abstract

Respiratory syncytial virus (RSV) is a common cause of respiratory infections, causing significant morbidity and mortality, especially in young children. Why RSV infection in children is more severe as compared to healthy adults is not fully understood. In the present study, we infect both pediatric and adult human nose organoid-air liquid interface (HNO-ALIs) cell lines with two contemporary RSV isolates and demonstrate how they differ in virus replication, induction of the epithelial cytokine response, cell injury, and remodeling. Pediatric HNO-ALIs were more susceptible to early RSV replication, elicited a greater overall cytokine response, demonstrated enhanced mucous production, and manifested greater cellular damage compared to their adult counterparts. Adult HNO-ALIs displayed enhanced mucus production and robust cytokine response that was well controlled by superior regulatory cytokine response and possibly resulted in lower cellular damage than in pediatric lines. Taken together, our data suggest substantial differences in how pediatric and adult upper respiratory tract epithelium responds to RSV infection. These differences in epithelial cellular response can lead to poor mucociliary clearance and predispose infants to a worse respiratory outcome of RSV infection.

## INTRODUCTION

Respiratory tract infections (RTIs) are a common cause of global mortality in children younger than 5 years old. Infection with respiratory syncytial virus (RSV) is the leading cause of infant deaths, resulting in approximately 100,000 to 200,000 deaths per year, predominantly in low-resource countries (1). RSV infection is the most common cause of bronchiolitis and pneumonia in children, which can clinically manifest as increased work of breathing caused by either partial or complete obstruction of the small distal airways by cellular debris and mucous (2–4). The epidemiologic burden of RSV is high; nearly every child has been infected prior to age 2, and reinfection is common throughout life (5, 6). Globally, RSV infection hospitalizes over 3 million children under 5 years old each year (1). While healthy adults infected with RSV predominantly have mild, self-limited RTIs, elderly adults are also at significant risk of morbidity and mortality (7). Currently, the treatment of RSV infection is largely supportive, and there are no approved anti-viral therapies other than aerosolized ribavirin (8) that is sparingly used due to potential toxicity and equivocal benefit.

The understanding of RSV pathogenesis and potential novel therapeutics have been slowed by the lack of pre-clinical models that recapitulate RSV infection in infants and elderly adults. Recently, using a non-invasive collection of stem cells from the nasal epithelium, we described the development of human nose organoid (HNO) cell lines that differentiated into a pseudostratified epithelium in an air-liquid interface (ALI) culture (9). The cell composition of the HNO-ALI culture mimics the epithelium of the upper respiratory track (9, 10). HNOs consist of several key cell types including 1) mucous-producing goblet cells, 2) apical ciliated cells, which function to move mucous, particulate matter, and pathogens out of the airway, 3) basal cells which function as the stem cells of the airway, and 4) club (secretory) cells, which secrete antiviral compounds (11). Apical ciliated cells of the HNOs, are the major cell type permissive to RSV infection, and when compared to clinical data, the epithelium demonstrates similar cytokine response to infection as seen in pediatric RSV infection (9, 12). Finally, a single organoid cell line reflects the genetic background of one individual, and thus HNOs from different individuals can be used to understand viral pathogenesis in vulnerable populations, such as young children.

Populations that are at higher risk of severe clinical RSV infection include premature infants, infants under 6 months of age, those with existing cardiopulmonary disease, the immunocompromised, and the elderly (13–15). In children, the largest group needing hospitalization are previously healthy, young infants (16). The increased hospitalization is thought to be due to smaller airway size which can result in a higher likelihood of lower distal airway obstruction and thereby respiratory distress (17). Alternatively, most healthy adults with RSV only have mild upper respiratory symptoms (18). A major gap in knowledge is the contribution of the respiratory epithelium to disease outcome. The epithelial cytokine response of the upper respiratory tract (URT) during RSV infection orchestrates the early innate and subsequent adaptive immune response. A robust immune response can play an important role in the control of viral replication and prevent dissemination into the lower respiratory tract (LRT) (19). It is unknown whether inability of the infant’s URT epithelium to control RSV infection compared to that of the adult can lead to increased susceptibility to RSV lower respiratory tract infection (LRTI). The HNOs derived from pediatric and adult donors provided us a unique opportunity to study the role of the nasal respiratory epithelium in response to RSV infection and disease outcome.

We characterized the viral kinetics, innate epithelial cytokine responses, and cell injury and remodeling to infection in both pediatric and adult derived HNOs with two contemporaneous RSV strains (RSV/A/Ontario [ON] and RSV/B/Buenos Aires [BA]). We found that pediatric-derived HNO-ALIs had more rapid RSV replication kinetics, higher overall cytokine responses, and greater loss of apical ciliated epithelium. Pediatric-derived HNO-ALIs had increased mucous area at baseline; however, both adult and pediatric HNO-ALIs demonstrated increased mucous production during RSV infection. Lastly, RSV/A/ON induced a greater cytokine response while RSV/B/BA infection resulted in greater cell damage in both adult and pediatric-derived HNO-ALIs. Rapid viral growth, dysregulated cytokine response, increased cellular damage, and an impaired mucociliary clearance in pediatric HNO-ALIs are all hallmarks of RSV disease in children. Taken together, these data suggest an epithelial-driven mechanism for predisposition of infants to severe LRTI in children.

## RESULTS

### Replication kinetics and morphologic analysis of RSV infected pediatric HNO-ALIs was distinct as compared to adult HNO-ALIs

To determine if there are differences in viral kinetics between pediatric and adult HNO-ALIs, four pediatric and four adult-derived HNO-ALIs were infected with RSVA/ON and RSVB/BA. Both RSV/A/ON and RSV/B/BA had similar viral concentration at peak infection (6 to 8 × 10^7^ viral RNA copies/mL). In pediatric HNO-ALIs, peak virus gene copy number occurred by day 2 with steady state achieved by 5 dpi, while in adult HNO-ALIs, peak infection occurred later at 5 dpi with steady state achieved by 8 dpi (Figure 1A, C; Figure S1 A, C, E & G). On the basolateral side, there were low levels of viral RNA detection at 5 and 8 dpi (approximately 1.0 × 10^7^ viral copies/mL), but never reached the level detected on the apical side (data not shown).

**Figure 1.**
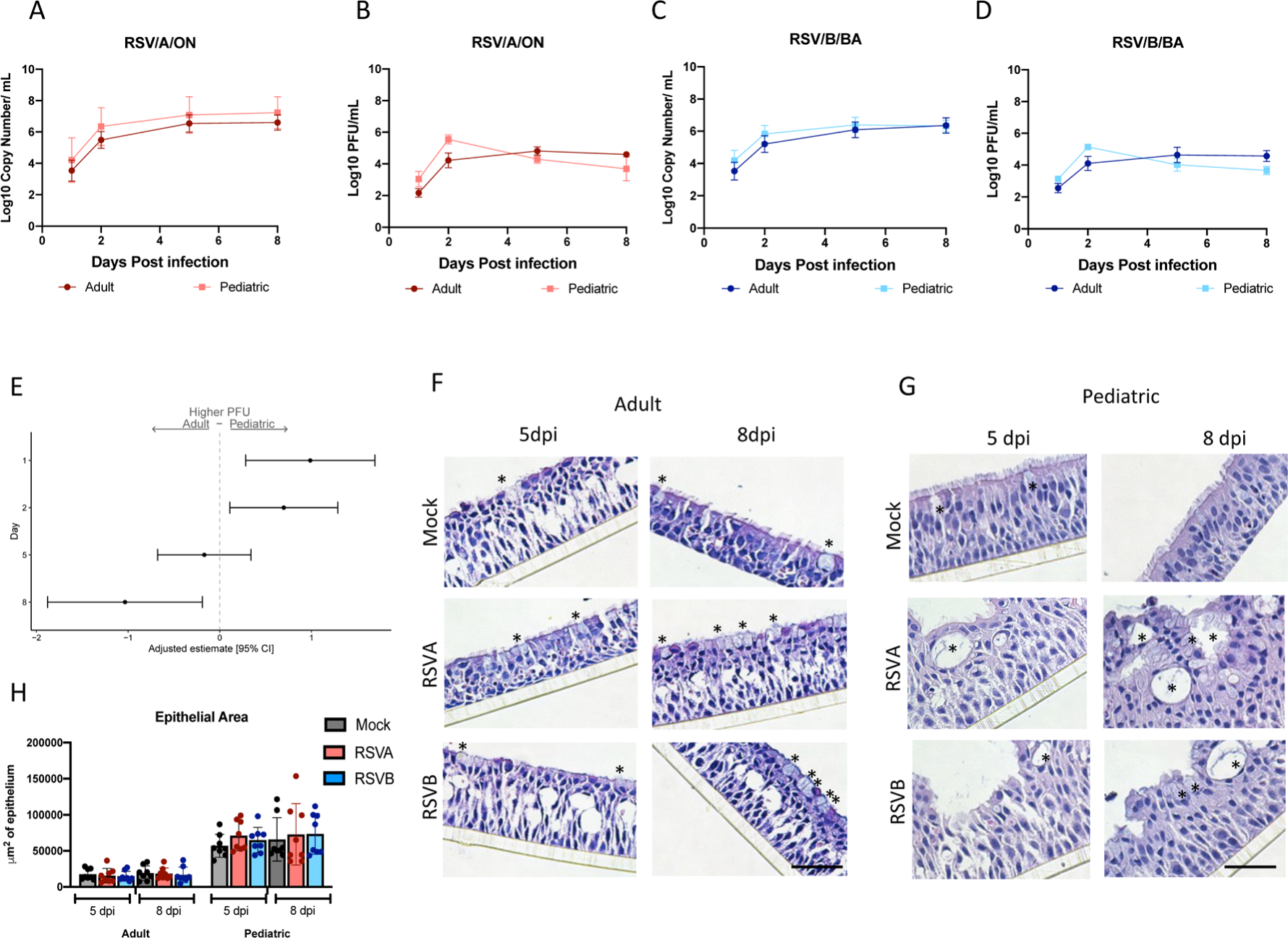
Replication kinetics and morphologic analysis of RSV infected pediatric HNO-ALIs and adult HNO-ALIs. **A)** Apical RT PCR for 4 adult and 4 pediatric HNO-ALIs for RSV/A/ON and RSV/B/BA and **B)** corresponding plaque data. **C)** Apical RT PCR for 4 adult and 4 pediatric HNO-ALIs for RSV/B/BA and **D)** corresponding plaque data. **E)** PFU data was modeled using the covariates: HNO age interacting with dpi, and viral infection. Forest plot demonstrating the adjusted beta coefficient estimate with 95% confidence intervals, represented by dots and T bars, respectively, between adult and pediatric HNO-ALIs. **F**) H&E images of adult HNO-ALI 918 at 5 and 8 day post-infection (dpi). (* denotes goblet cells). **G**) H&E images of pediatric HNO-ALI 9003 at 5 and 8 dpi. (* denotes goblet cells). **H**) Comparison of epithelial area between pediatric and adult HNO-ALIs at 5 and 8 dpi with RSV/A/ON, RSV/B/BA, or mock infection. Scale bar is 100 µm.

Infectious virus by plaque assay showed similar kinetics to the viral RNA. Interestingly pediatric HNO-ALIs demonstrated a peak viral titer at 2 dpi which declined at the later time points, whereas in adult HNO-ALIs, RSV viral titer peaked at 5 dpi and achieved steady state at 8 dpi (Figure 1B, D; Figure S1 B, D, F & H). Live infectious virus was not detected on the basolateral compartment suggesting that low amounts of viral RNA were not infectious (data not shown).

To determine if there were significant differences in the RSV replication kinetics, we utilized a linear regression analysis to compare RSV replication in pediatric versus adult HNO-ALIs. In comparing viral copy numbers, pediatric HNO-ALIs had a statistically significant higher viral copy number (p=0.048) than adult HNO-ALIs, across all time points (adjusted mean difference: 0.54, 95% CI: 0.01, 1.08) whether infected with RSV/A/ON or RSV/B/BA. Interestingly, pediatric HNO-ALIs had a significantly higher number of infectious virions measured by a quantitative plaque assay at 1 and 2 dpi (adjusted mean difference at 1 dpi: 0.99, 95% CI: 0.28, 1.69; at 2 dpi: 0.70, 95% CI: 0.11, 1.29). However, by 8 dpi the live virus titers in pediatric HNO-ALIs were significantly lower than adults suggesting the virus in pediatric HNO-ALIs replicated robustly early in the infection, but later into the infection the virus was not replicating as efficiently (Figure 1E).

H&E analysis of pediatric and adult-derived HNO-ALIs demonstrated robust changes in cellular morphology compared to mock infection. There was a loss of apical ciliated cells and expansion of goblet cells especially at 5 and 8 dpi in both adult and pediatric HNOs (Figure 1F-G). As compared to adult HNO-ALIs, Pediatric HNO-ALIs displayed greater morphological changes such as thicker epithelium, with increased large vacuole-like formations, some of which via immunostaining contained mucus and/or cilia (as shown by immunolabeling, Figure 1F-G). We measured the overall epithelial area of both adult and pediatric HNO-ALIs and did not detect a difference with regards to changes with viral infection or with time (Figure S1 I and J). However pediatric HNO-ALIs had a larger total area of epithelium compared to adult HNO-ALIs at baseline (aRR: 3.56, 95% CI 2.98-4.25). There was no statistically significant difference in epithelial area across the four adult and four pediatric HNO-ALIs lines with regards to viral infection or time (Figure 1H), although there was variability among different HNO-ALIs lines. HNO2 showed a trend toward overall decreased epithelial area with RSV infection (Figure S1I). Overall, viral copy numbers in RSV infected pediatric HNO-ALIs were detected at higher levels throughout the course of infection, whereas infectious virus peaked earlier and decreased more rapidly as compared to kinetics of adult HNO-ALI. Although the overall area of the epithelium was thicker in pediatric HNO-ALI lines, this was not affected by active virus replication.

### Epithelial cytokine and chemokine response during RSV infection in pediatric and adult HNO-ALIs

To characterize host-specific changes in cytokine secretion during RSV infection, we performed a 24 cytokine Luminex analysis at 1, 2, 5, and 8 dpi (supplementary data Set 1). In general, cytokine levels peaked when virus titers peaked, and subsequent steady state was achieved. Thus, in pediatric HNO-ALIs levels began to rise by 2 dpi, peaking at 5 and 8 dpi (Figure S2), and in adult HNO-ALIs this occurred later at 5 and 8 dpi (Figure S3). In both adult and pediatric HNO-ALIs, cytokines classically induced in human RSV infection such as RANTES, IP-10, IL-6, and IL-8, were induced in both the basolateral and apical compartments after infection with RSV/A/ON and RSV/B/BA (Figures S2, S3).

To further understand the cytokine secretion in response to RSV infection, we analyzed the overall cytokine expression in terms of composite Z-score probabilities to determine if there were significant differences in the epithelial cytokine response both at baseline and during infection with RSV between the pediatric and the adult HNO-ALIs. Un-infected pediatric HNO-ALIs demonstrated a trend toward higher overall cytokine expression as compared to adults (aOR: 1.23; 95% CI: 0.97-1.555, Figure 2A). After infection with RSV, pediatric HNO-ALIs had a significantly higher overall cytokine expression in both RSV/A/ON (aOR: 1.69; 95% CI: 1.33-2.14) and RSV/B/BA (aOR: 1.78;95% CI: 1.41-2.25) infected HNO-ALIs as compared to adults. We compared levels between pediatric and adult HNO-ALIs (Figure 2A) after grouping the cytokines by their biological function, which included anti-viral (IL-29), chemoattractant (MCP-1, MCP-3, MIP-1α, MIP-1β, MIG/CXCL-9, IP-10, eotaxin, RANTES), maturational (VEGF-α, FGF-2, G-CSF, GM-CSF), metalloproteinase and its inhibitor (MMP-7, MMP-9, TIMP), pro-inflammatory (IL-1α, IL-1β, TNFα, IL-6, IL-8) and regulatory cytokines (TGF-β, IL-17E). Interestingly, while nearly every functional cytokine groups had significantly higher overall expression in pediatric HNO-ALIs infected with RSV, a notable exception was the regulatory cytokine group, which trended lower in pediatric HNO-ALIs compared to adult HNO-ALIs (RSV/A/ON aOR: 0.70; 95%: CI: 0.48-1.04, & RSV/B/BA aOR: 0.82; 95% CI: 0.56-1.20) (Figure 2A).

**Figure 2.**
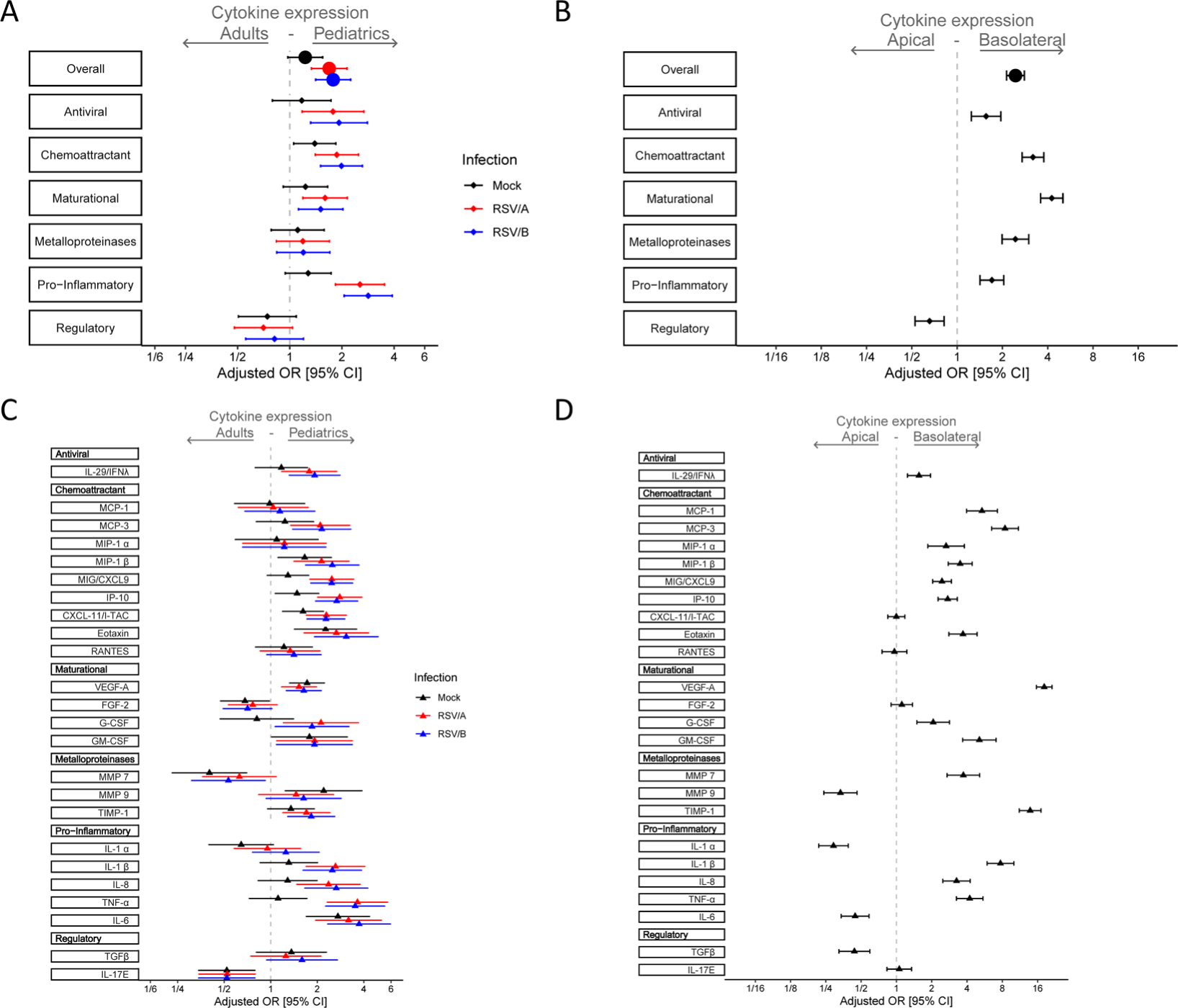
Epithelial cytokine and chemokine response during RSV infection in pediatric and adult HNO-ALIs. Cytokine/chemokine secretion was modeled on the following factors: HNO age interacting with viral infection, dpi interacting with RSV, and cell surface. Adjusted odds ratio estimates and their associated 95% confidence intervals are represented by dots and T-bars, respectively. Mock infection is shown in black, RSV/A/ON in red, and RSV/B/BA in blue. **A)** Forest plot showing the adjusted odds ratio with 95% confidence interval for overall cytokine secretion and cytokine group expression between adult and pediatric HNO-ALIs. **B)** Forest plot showing the adjusted odds ratio with 95% confidence interval cytokine expression for overall cytokine secretion and cytokine group expression in the apical or basolateral compartment. **C)** Forest plot of adjusted odds ratio with 95% confidence intervals for individual cytokines in each cytokine group for mock, RSV/A/ON, and RSV/B/BA infection between adult and pediatric HNO-ALIs. **D)** Forest plot of adjusted odds ratios with 95% confidence intervals for individual cytokines in the apical or basolateral compartment.

To explore our model further, we examined apical vs. basolateral secretion of cytokines, which would mimic the cytokines released into the airway versus into the circulation, respectively (Figure 2B). We found that in general the basolateral compartment had overall higher secretion of all cytokines (aOR: 2.44; 95% CI: 2.13-2.80). Intriguingly, chemoattractants like MCP and eotaxin, and maturational factors such as VEGF-α were predominantly found in the basolateral compartment (aOR: 18.21; 95% CI: 15.63-21.22), which was consistent with their physiologic role in recruitment of immune cells and in angiogenesis.

We next looked at individual cytokines between pediatric and adult HNO-ALIs and their location (Figure 2C and 2D). IFN-lambda/ IL29, a type III interferon essential for anti-viral signaling, was detected at higher levels in the RSV infected pediatric HNO-ALIs (Figure 2C) and predominantly in the basolateral compartment where active virus replication was not occurring (Figure 2D). Chemokines IP10, CXCL11, CXCL9 that are involved in the recruitment and maturation of the immune response, were highly upregulated in pediatric HNO-ALIs after infection with RSV/A/ON and RSV/B/BA (Figure 2C), plus IP-10 and CXCL9 were detected at significantly higher concentration in the basolateral compartment (Figure 2D). Expression of maturational factors FGF-2 and GM-CSF did not change significantly with infection or time; however, G-CSF was increased with infection in the pediatric HNO-ALIs compared to adults. There was no appreciable difference in the expression of matrix metalloproteinases (MMP7 and MMP9) during infection with RSV, potentially due to the presence of TIMP (inhibitor of MMPs), which increased at modest levels at 5 and 8 dpi. Proinflammatory cytokines, in general, were higher in RSV infected pediatric HNO-ALIs and found at higher concentrations in the basolateral compartment. TNFα, a pro-inflammatory cytokine involved with cell necrosis, was also highly induced in pediatric compared to adult HNO-ALIs after infection with RSV. The two regulatory cytokines that we measured, TGF-β and IL-17E, did not increase significantly with RSV infection, but differed in their expression pattern. IL-17E was detected at significantly higher levels in the adult HNO-ALIs, while TGF-β was seen at higher concentrations in the pediatric HNO-ALIs, although not statistically significant and mostly in the apical compartment. Lastly, there were notable exceptions in the predominant location of several cytokines, IL-1α, MMP9, IL-6, and TGFβ, which favored the apical compartment, demonstrating differences in the location cytokines are released with a minority of cytokines we studied favoring the apical compartment (consistent with airway) where virus infection is occurring versus the majority of cytokines favoring the release into the basolateral compartment (consistent with circulation) (Figure 2D).

Taken together, this suggests that pediatric HNO-ALIs infected with RSV elicited a higher overall cytokine response, predominantly in the pro-inflammatory and chemoattractant groups, that can result in a heightened activation of the immune system. This heightened response is less likely to be moderated by regulatory cytokines, which were expressed in lower amounts in pediatric HNO-ALIs compared to adults.

### Cell death and damage during RSV infection in pediatric and adult HNO-ALIs

We sought to quantify cellular damage and death via LDH and caspase 3/7 activity which measure cellular injury and apoptosis, respectively, during RSV infection. We postulated levels would be higher at the site of active RSV replication, the apical compartment. Overall, there was a low baseline secretion of LDH in the apical and basolateral compartments, but after infection with RSV, there was a notable increase that peaked at 5 and 8 dpi predominantly in the apical compartment (Figure 3A-C and Figure S4A, C, E & G) with minimal change in the basolateral compartment (Figure S5A, C, E & G). Caspase 3/7 activity paralleled those of LDH, with overall a low level of apoptosis detected in mock infections and increased activity after either RSV/A/ON or RSV/B/BA infection at 5 and 8 dpi only in the apical surface (Figure 3D-F, Figure S4 B, D, F & H). A statistically significant correlation (r = 0.832; p<0.002) for adult HNO-ALIs, and (r=0.625; p=<0.002) for pediatric HNO-ALIs, was observed between LDH and caspase activity (Figure 3G). Interestingly, RSV infected pediatric HNO-ALIs had higher LDH activity relative to caspase than adult suggesting higher non-apoptotic cellular damage as compared to adults.

**Figure 3.**
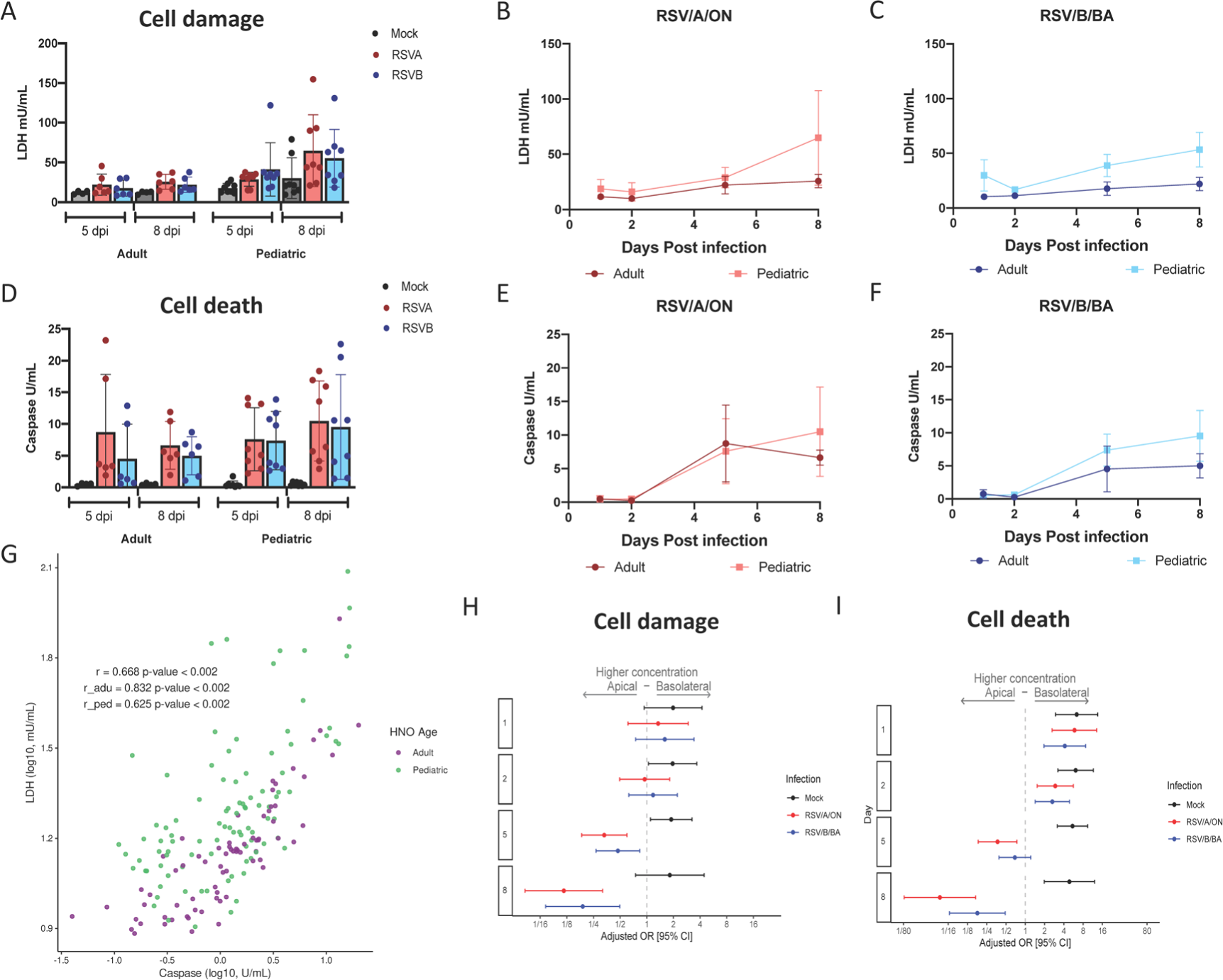
Cell death and damage during RSV infection in pediatric and adult HNO-ALIs. LDH and caspase area were modeled on the following factors: interacting cell surface, viral infection, and dpi as well as age. **A)** Amount of cell damage, measured by apical LDH, at day 5 and day 8 in adult vs pediatric HNO-ALIs, infected with RSV/A/ON, RSV/B/BA, or mock infection. **B)** Amount of apical LDH in 4 pediatric versus 4 adult HNO-ALIs infected with RSV/A/ON. **C)** or RSV/B/BA. **D)** Amount of cell death, measured by apical caspase, at day 5 and day 8 in adult vs pediatric HNO-ALIs, infected with RSV/A/ON, RSV/B/BA, or mock infection. **E)** Amount of apical caspase in 4 pediatric vs 4 adult HNO-ALIs infected with RSV/A/ON. **F)** or RSV/B/BA. **G)** Spearman correlation of pediatric and adult levels of LDH and caspase. Each dot represents the mean value of each unique experimental condition where both LDH and caspase were measured. Adult and Pediatric samples are colored magenta and green, respectfully **H)** Forest plot with adjusted odds ratio with 95% confidence intervals of apical or basolateral expression of LDH. **I**) Forest plot of adjusted odds ratios of apical or basolateral expression of caspase. Adjusted odds ratio estimates with 95% confidence intervals are represented by dots and T-bars, respectively.

A multivariable analysis was used to assess differences between cell damage (LDH) and cell death (caspase) after infection with RSV/A/ON and RSV/B/BA. LDH level in the apical compartment was significantly higher in pediatric compared to adult HNO-ALIs (aOR 3.89; 95% CI: 2.44-6.18), while levels of apoptosis (caspase 3/7) was comparable between the two groups (aOR 1.04; 95% CI: 0.77-1.44). However, when examining three-way interactions between cell surface (apical vs basolateral), infection (mock, RSV/A/ON, RSV/B/BA) and time, we found that while mock infection favors the basolateral compartment for expression of both LDH and caspase, as infection with both RSV/A/ON and RSV/B/BA progresses, detection increased significantly at 5 and 8 dpi to favor apical expression of LDH and caspase (Figure 4H-I). This data suggests that while there are low levels of basal cell turnover in HNO-ALIs, infection with RSV causes a predominantly apical increase in cell damage and death, consistent with localization of the virus detected by qRT-PCR and plaque assays. In addition, apical cellular injury from a non-apoptotic pathway is more prominent in pediatric HNO-ALIs compared to adults in response to RSV infection.

**Figure 4.**
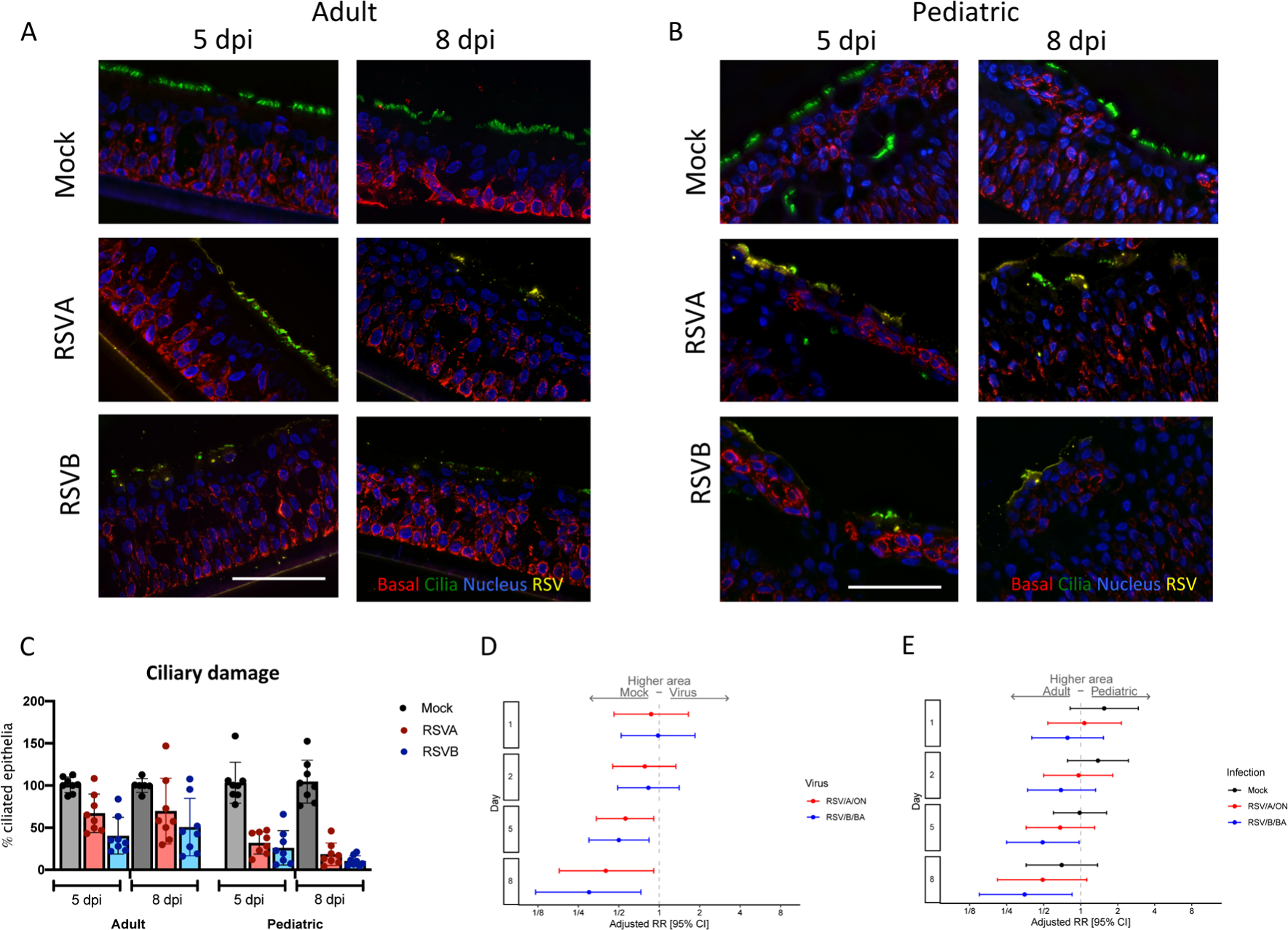
Ciliary damage in pediatric and adult HNO-ALIs. **A)** Representative IF imaging of a single adult HNO at 5 and 8 days post-infection (dpi). Basal cells are stained in red by Krt5, ciliated cells are stained in green by acetylated alpha tubulin, RSV particles are stained in yellow by anti-RSV antibody, and cellular nuclei are stained in blue by DAPI and **B)** single pediatric HNO-ALI at 5 and 8 dpi. Basal cells are stained in red by Krt5, ciliated cells are stained in green by acetylated alpha tubulin, RSV particles are stained in yellow by anti-RSV antibody, and cellular nuclei are stained in blue by DAPI. Scale bar is 100 µm. **C)** Percentage of ciliary damage in adult vs pediatric HNO-ALIs at 5 or 8 dpi. In this graph, the ciliary area is normalized to the corresponding HNO-ALI mock infection being 100%. **D)** Forest plot of mock versus virus of cilia area modeled on the following factors: the interaction of dpi and age, the interaction of age and viral infection, and the interaction of dpi and virus. Adjusted risk ratio estimates and their associated 95% confidence intervals are represented by dots and T-bars, respectively. **E)** Forest plot of ciliary area of adults versus pediatrics was modeled on the following factors: the interaction of dpi and HNO-ALI age, the interaction of age and viral infection, and the interaction of dpi and virus. Adjusted risk ratio estimates and their associated 95% confidence intervals are represented by dots and T-bars, respectively.

### Pediatric HNO-ALIs had higher amount of ciliary damage compared to adult HNO-ALIs with RSV infection

Immunofluorescence image quantification for apical ciliated cells, goblet cells, secretory cells, basal cells, and RSV localization was performed to examine cell-type specific changes of HNO-ALIs during RSV infection. Because RSV infected pediatric HNO-ALIs experienced greater release of proinflammatory cytokines and higher biomarker of cell damage compared to adults, we predicted greater ciliary cell damage. As expected, RSV infection was localized only to the apical ciliated cells (Figure 4A-B). Using fluorescence thresholding normalized to mock, ciliated area (quantified by acetylated tubulin or ace-tub) was unaffected during early infection but decreased during late infection (5 or 8 dpi) (Figure 4C). In pediatric HNO-ALIs, there was a nearly total loss of ciliated epithelium demonstrated by loss of acetylated tubulin immunolabeling (Figure 4B-C).

Linear regression models that included age (adult or pediatric), time of infection (day 1, 2, 5, and 8), virus (mock, RSV/A/ON and RSV/B/BA) with their interaction terms (day-virus, age-virus, and day-age) were generated to determine if age, virus or time had an impact on the cell composition. We compared apical ciliary damage between adult and pediatric HNO-ALIs. Because each HNO-ALIs has a different amount of ciliary area at baseline, we normalized the ciliary area to their respective mock HNO-ALI line (i.e. 100%). While both adult and pediatric HNO-ALIs had decreased apical ciliary area during RSV infection, pediatric HNO-ALI lines had approximately three to five times more ciliary damage when compared to adult lines at 8 dpi with RSV/A/ON and RSV/B/BA respectively. (Figure 4C, Figure S6A-B).

We then analyzed the total area of cilia (not normalized to mock) and generated a relative risk of apical ciliary damage compared to mock infection. There was no significant difference between viral infection and mock at early timepoints, 1 and 2 dpi. However, 5 and 8 dpi, RSV/A/ON and RSV/B/BA resulted in significantly reduced apical ciliary area as compared to mock (RSV/A/ON: 5 dpi aRR: 0.56; 95% CI: 0.34-0.91; 8 dpi aRR: 0.40; 95% CI: 0.18-0.90; and RSV/B/BA: 5 dpi aRR: 0.50; 95% CI: 0.30-0.84; 8 dpi aRR: 0.30; 95% CI: 0.12-0.73) (Figure 4D). Interestingly, pediatric HNO-ALI lines infected with RSV/B/BA at 5 and 8 dpi had significantly less total area of apical ciliated cells as compared to adult lines (5 dpi aRR: 0.49; 95% CI: 0.25-0.97; 8 dpi aRR: 0.35; 95% CI: 0.15-0.85). Late infection with RSV/A/ON in pediatric compared to adult HNO-ALIs had a trend towards increased loss of apical ciliary area (8 dpi aRR: 0.49; 95% CI: 0.12-1.12). Taken together both RSV/A/ON and RSV/B/BA induced significant apical ciliary cell destruction that peaked at 8 dpi with greater damage visualized in pediatric HNO-ALIs compared to adults. The ciliary damage was greater with RSV/B/BA than RSV/A/ON.

### Pediatric HNO-ALIs had higher baseline levels of goblet cells and produced larger amounts of mucous compared to adult HNO-ALIs

Increase mucous production is a hallmark of RSV infection in children and adults. We hypothesized that RSV infection would induce an increase in goblet cells and mucous production in both pediatric and adult HNO-ALIs. Mucous-producing goblet cells were measured by staining with Muc5AC. Two parameters were evaluated, the number of mucous-producing goblet cells and the amount of area staining for mucous. Pediatric HNO-ALIs had a larger, more diffuse staining of Muc5AC prior to infection compared to adults (Figure 5A-C), while hypersecretion of mucus was noted during late infection (5 and 8 dpi) (Figure 5C-D, Figure S6C-D). Increase in percentage of mucous producing goblet cells was more apparent after infection with RSV/B/BA than by RSV/A/ON (Figure 5E-F) in both adults and pediatric HNO-ALIs. When examining overall mucus production, we observed that while at baseline adult and pediatric HNO-ALIs had similar percentage of goblet cells (Muc5ac staining with DAPI+ nuclei/ total number of DAPI+ cells), these same percentage of cells in pediatric HNO-ALIs produced nearly twice as much mucous (area of Muc5ac staining, Figure 5A-C) even prior to infection with RSV (1 dpi aRR: 2.44, 95% CI: 1.64-3.64). In early infection with RSV (1 and 2 dpi), pediatric HNO-ALIs had a significant increase in mucus area compared to adult HNO-ALIs (Figure 5B). As infection progressed (5 and 8 dpi), the difference between mucus production in adult and pediatric HNO-ALI decreased, likely due to both pediatric and adult increasing overall mucus production at late infection (Figure 5B-C). Similarly, when examining how mucus area changed over time compared to mock infections, we found at 5 and 8 dpi, both the area of mucous and percentage of mucous producing goblet cells increased, with infection with RSV/B/BA causing a higher increase in the percentage of mucous producing goblet cells than in RSV/A/ON (Figure 5D, F), although not significantly different.

**Figure 5.**
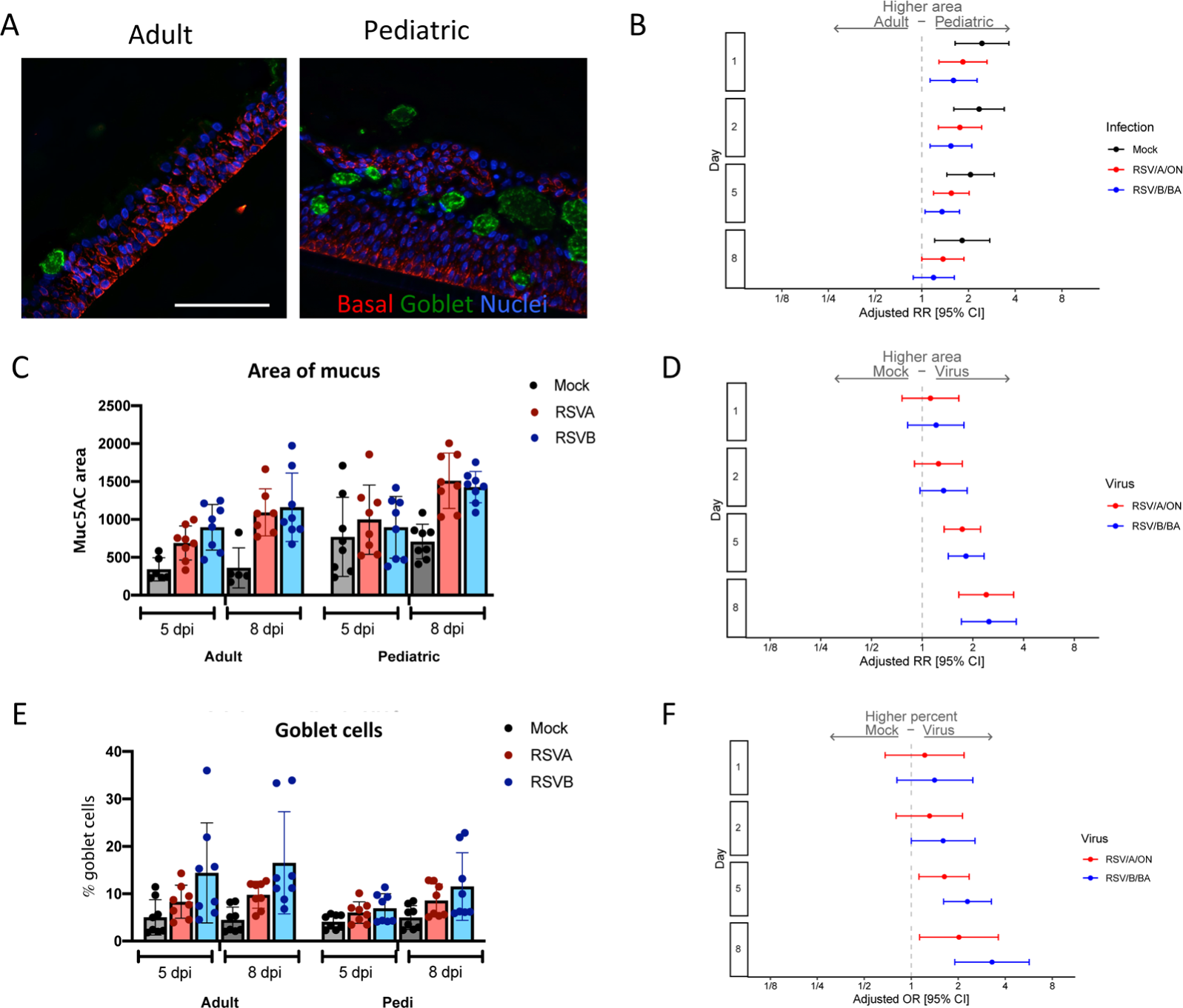
Mucous secretion in pediatric and adult HNO-ALIs. **A)** Representative IF imaging of a single uninfected adult and pediatric HNO-ALI. Basal cells are stained in red by Krt5, mucus in green by Muc5AC, and cellular nuclei are stained in blue by DAPI. **B)** Mucous area was modeled using the following factors: the interaction of days post-infection (dpi) and HNO age, the interaction of age and viral infection, and the interaction of dpi and virus. Forest plot of the adjusted relative risk with 95% confidence interval of having higher mucous area in pediatric vs adult HNO-ALIs. Adjusted risk ratio estimates and their associated 95% confidence intervals are represented by dots and T-bars, respectively. Scale bar is 100 µm. **C)** The area of mucus (Muc5AC+ area) in adult compared to pediatric HNO-ALIs infected with RSV/A/ON (red) and RSV/B/BA (blue) at 5 and 8 dpi. **D)** Adjusted relative risk of higher mucus (Muc5AC) area in mock compared to viral infection in combined adult and pediatric HNOs. **E)** Percentage of goblet cells (Muc5AC+ cells with DAPI+ nuclei over total number of DAPI+ cells) in adult compared to pediatric HNO-ALIs at 5 and 8 dpi. **F**) Goblet cell percentage data was modeled using the following factors: the interaction of age and virus as well as the interaction of day and virus. Adjusted odds ratio estimates and their associated 95% confidence intervals are represented by dots and T-bars, respectively.

### Secretory cells decreased in pediatric HNO-ALIs, but basal cells remained constant in both adult and pediatric HNO-ALIs

Club or secretory cells possibly play an anti-inflammatory role by their secretion of uteroglobin (also known as secretoglobin family 1A member 1 -SCGB1A1), and function as progenitor cells for themselves and apical ciliated cells. We postulated an increase in secretory cells late in RSV infection due to the significant ciliated cell damage caused by RSV. Secretory cells, measured by club cell frequency, accounted for a small population of cells in both adult and pediatric HNO-ALIs. We analyzed the percentage of club cells (number of CC10 positive cells/ DAPI+ nuclei) in pediatric and adult HNO-ALI lines after infection with RSV. Interestingly, while the percentage of club cells stayed relatively constant in adult HNO-ALIs, at 5 dpi with RSV/A/ON and RSV/B/BA, pediatric HNO-ALIs had reduced percentage of club cells compared to adult HNO-ALIs (RSV/A/ON aOR: 0.37; 95% CI: 0.20-0.69; & RSV/B/BA aOR: 0.38; 95% CI: 0.21-0.69) (Figure 6C, E). Moreover, at day 8, mock pediatric HNOs had significantly lower percentage of club cells compared to mock adult HNO-ALI lines. This difference between pediatric and adult lines in the mock HNO-ALIs is likely driven by an increase in club cells in a single adult HNO-ALI line (HNO2) rather than a true loss in the pediatric HNO-ALI lines, as club cells in mock pediatric HNO-ALIs were relatively stable over time (Figure S6E-F). Decrease percentage of club cells in RSV infected pediatric HNO-ALIs compared to adults may contribute to the enhanced proinflammatory response and cellular injury being observed in the pediatric HNO-ALIs infected with RSV.

**Figure 6.**
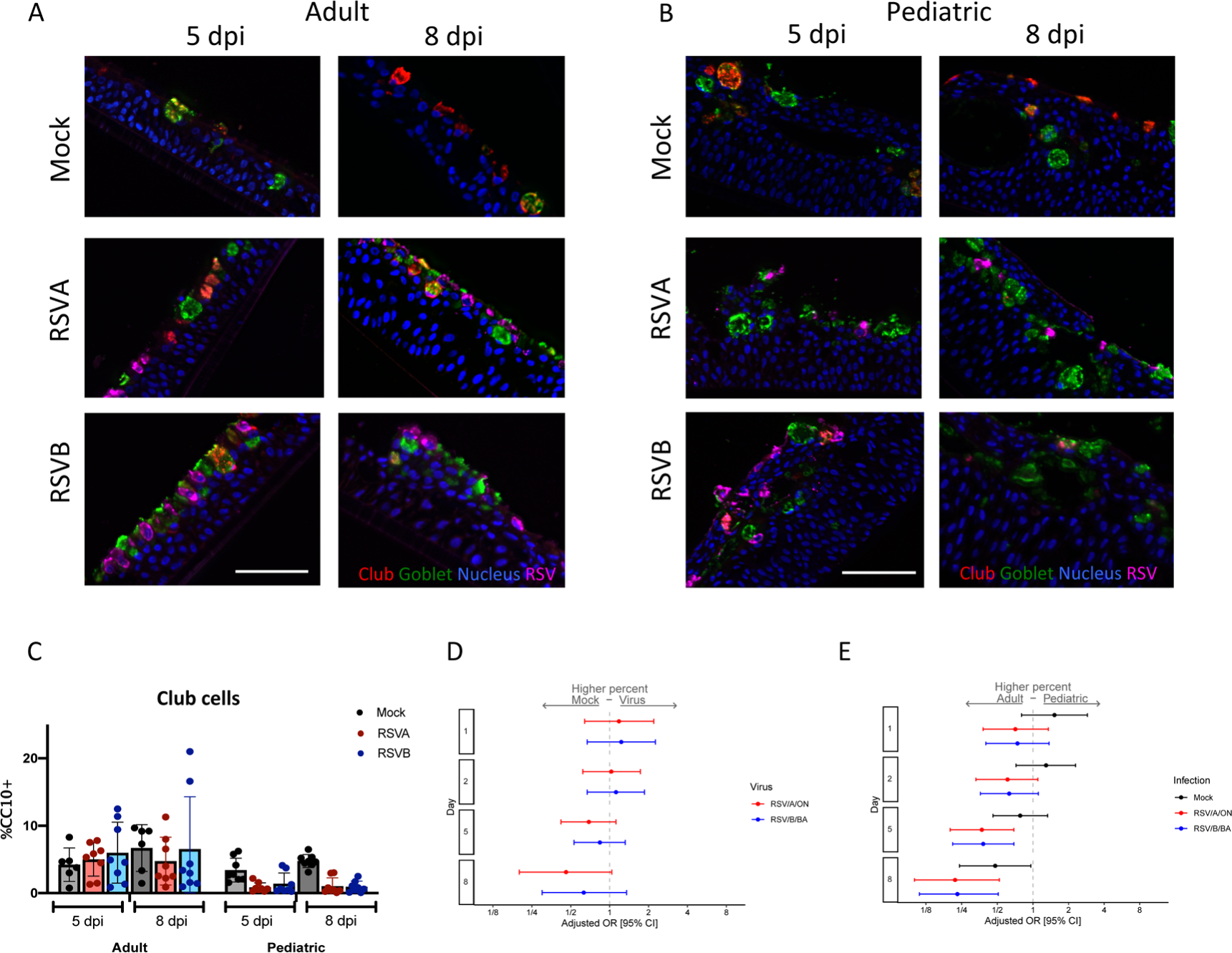
Club cells in pediatric and adult HNO-ALIs. **A)** Representative IF images of a single adult HNO and **B)** single pediatric HNO at day 5 and 8 post infection. Club cells are stained red with CC10, goblet cells are stained in green by Muc5AC, RSV particles are stained in magenta by anti-RSV antibody, and cell nuclei are stained in blue by DAPI. Scale bar is 100 µm. **C)** Percentage of club cells (CC10+ cells with DAPI+ nuclei over total number of DAPI+ cells) in adult compared to pediatric HNO-ALIs at 5- and 8-day post-infection (dpi). **D)** Club cell percentage data was modeled using the following factors: the interaction of dpi and HNO age, the interaction of age and viral infection, and the interaction of dpi and virus. Adjusted odds ratio estimates with 95% confidence intervals are represented by dots and T-bars, respectively. **E**) Adjusted odds ratio with 95% confidence intervals of higher percentage of club cells in adult compared to pediatric HNOs.

We also measured percentage of basal cells (airway stem cells which can differentiate to replenish other lost cell population) using Krt5 staining (number of Krt5+ cells/ DAPI+ cells). Compared to adult HNO-ALIs, pediatric HNO-ALIs had nearly twice as many (39% vs 79%) Krt5+ basal cells (OR 5.49, 95% confidence interval 4.35-6.49). There was no significant difference in the percentage of basal cells after infection with RSV in neither pediatric nor adult HNO-ALIs (Figure S6G-H).

### Cell proliferation during RSV infection in adult and pediatric HNO-ALIs

We measured the level of cell proliferation during RSV infection in pediatric and adult HNO-ALIs by Ki67 staining. Both adult and pediatric HNO-ALIs had a low level of baseline proliferation as measured by percentage of Ki67 positivity (approximately 2-10%) (Figure 7A and B). However, there was significant variability between cell lines, with HNO-ALI 918 demonstrating a high 20% Ki67 positivity at baseline (Figure 7F). Baseline proliferation in mock infected pediatric HNO-ALIs was approximately 2-fold higher as compared to adult HNO-ALI lines at early timepoints (1 dpi aOR: 2.07, 95% CI: 1.19-3.62; 2 dpi aOR: 1.78, 95% CI: 1.12-2.84), however at 5 and 8 dpi they were not significantly different (Figure 7E). There was no statistically significant difference in the amount of cell proliferation associated with viral infection, although at 8 dpi there appears appreciable increase (Figure 5F-G).

**Figure 7:**
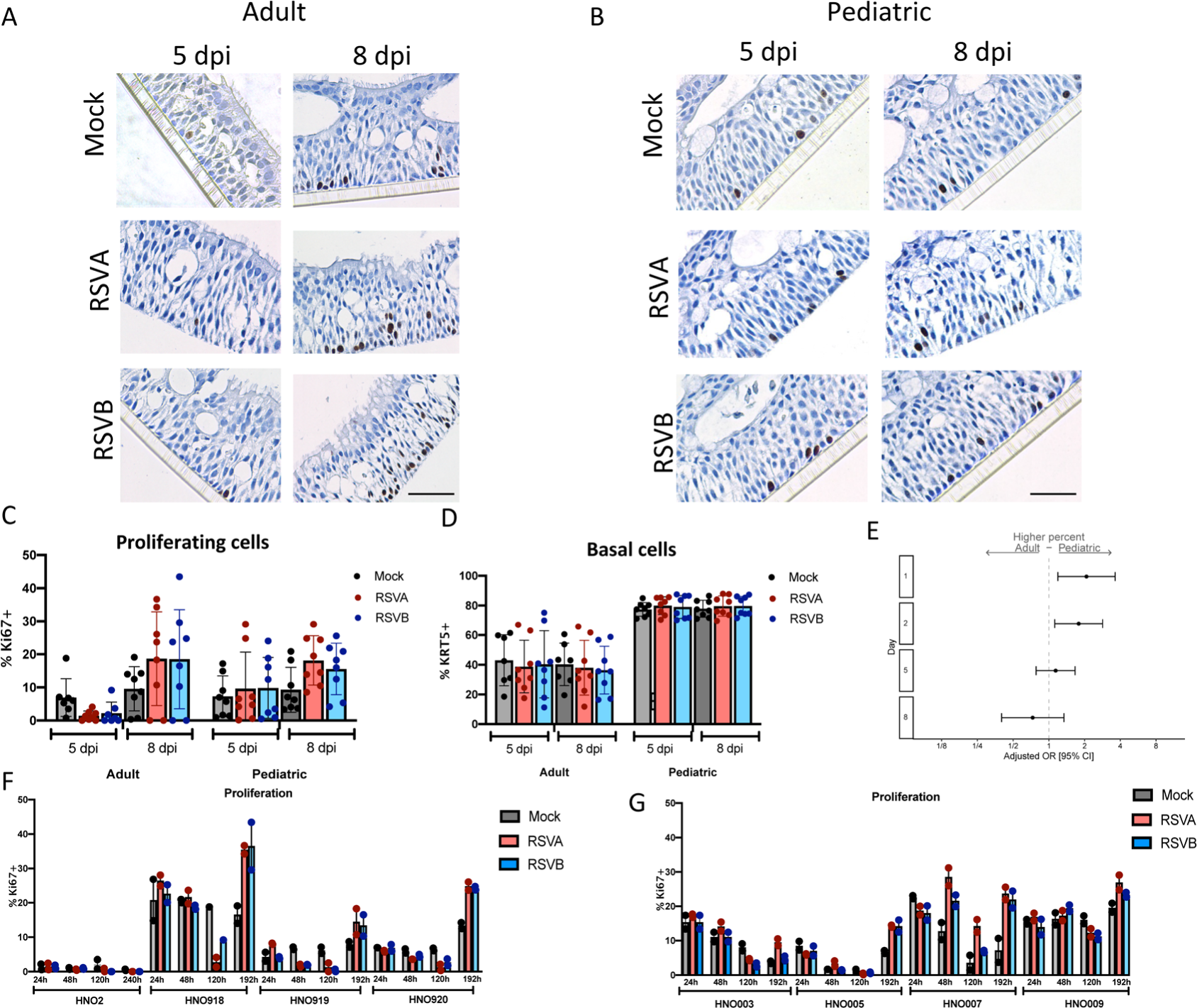
Cell proliferation in pediatric and adult HNO-ALIs. **A)** Representative Ki67 staining of a single adult HNO-ALI and **B)** single pediatric HNO-ALI at 5- and 8-day post infection (dpi) with RSV/A/ON, RSV/B/BA, or mock infection. Scale bar is 100 µm. **B)** Percentage of proliferating cells (number of Ki67 positive cells/ total cells) in 4 adult HNO-ALIs versus 4 pediatric HNO-ALIs at 5 or 8 dpi after infection with RSV/A/ON, RSV/B/BA, or mock infection. **D)** Percentage of basal cells (number of Krt5 positive cells/ total cells) in 4 adult HNO-ALIs versus 4 pediatric HNO-ALIs at 5 or 8 dpi after infection with RSV/A/ON, RSV/B/BA, or mock infection. **E)** Ki67 percent data was modeled using the following factors: the interaction of dpi and HNO age as well as the interaction of dpi and virus. Forest plot showing the adjusted odds ratio with 95% confidence intervals between adult and pediatric HNOs of having a higher amount of proliferating cells. Adjusted odds ratio estimates and their associated 95% confidence intervals are represented by dots and T-bars, respectively. There was no difference between mock and RSV infected HNOs, and thus this was dropped from the model and graph. **F)** Percentage of proliferating cells, in 4 adult HNO-ALIs at 1, 2, 5, and 8 dpi with RSV/A/ON or RSV/B/BA or mock infection. Each line shows variability in amount of proliferation. **G)** Percentage of proliferating cells in 4 pediatric HNO-ALIs at 1, 2, 5, and 8 dpi with RSV/A/ON or RSV/B/BA or mock infection.

## Discussion

RSV first enters and begins infection in the upper respiratory tract before it can extend into the lower respiratory tract. Therefore, understanding the role of the upper respiratory tract epithelium as the first line of defense against RSV infection may shed information on differences in outcomes between pediatric and adult RSV infection. We thus sought to examine differences in how the upper respiratory tract epithelium responds to RSV infection in young children and adults using an ex-vivo challenge model of the human nose organoid. We found that pediatric HNO-ALIs reached peak RSV titers as early as two days when compared to adults. Pediatric HNO-ALIs also elicited a greater cytokine response in almost all cytokine functional groups we tested except for regulatory cytokines. While both adult and pediatric HNO-ALIs demonstrated significant apical ciliary damage and enhanced mucous production, pediatric HNO-ALIs endured heightened ciliary damage, likely due to higher pro-inflammatory cytokines and possibly a reduction in club cells, a model summarizing these findings is depicted in Figure 8. Our findings mirrored to those found in young infants, who have airway occlusion due to mucus and cellular debris and relatively small airway size (17). This poor mucociliary clearance can impede viral elimination and lead to cell-to-cell spread or aspiration of infectious cellular debris causing LRTI.

**Figure 8:**
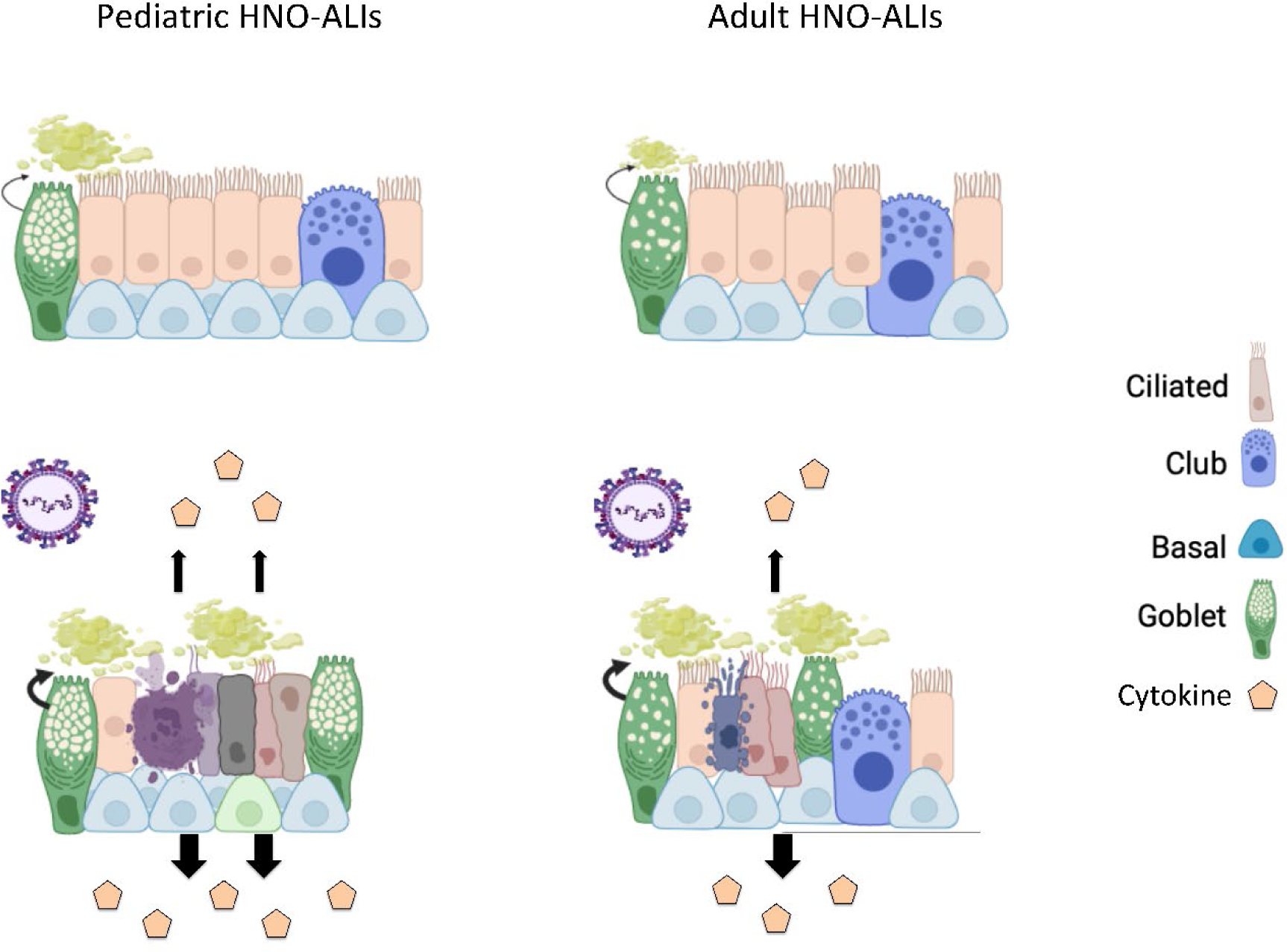
Schematic of RSV infection in pediatric compared to adult HNO-ALIs. Prior to infection, pediatric lines have increased mucus as well as more basal cells. After infection with RSV, both pediatric and adult HNOs have ciliary damage, increased mucus production, cell death and damage, and increase cytokine production. However, pediatric lines have more ciliary damage, more cellular damage, and increased cytokine production compared to adult lines. Pediatric lines do not have a comparative increase in the amount of apoptosis to their increased cell damage compared to adult lines, suggesting that other mechanisms of cell death such as cell necrosis may be contributing, further augmenting the inflammatory response. Created with BioRender.com

Each of our HNO-ALIs represents the genetic background of the individual it was derived from, and thus can be used to model differences in viral-host interaction throughout the population. Our pediatric HNO-ALIs were obtained from age ranges of 3 months to 1.5 years and the adults from 39-65 years of age, which provided a platform to compare adult and infant responses to RSV infection. Pediatric HNO-ALIs showed inherent characteristics that were distinct from that of adults; they were quick to propagate during cultivation of HNO-ALIs, had thicker epithelium and higher baseline levels of goblet cells. Pediatric HNO-ALIs were also more severely impacted with RSV infection as compared to adults. RSV replication as detected by viral RNA and infectious virions was faster and detected at higher levels in pediatric HNO-ALIs compared to adult HNO-ALIs. Pediatric HNO-ALIs evidenced approximately a 2-fold increase in apical LDH in late infection consistent with more cell damage of the apical ciliated cells, the site of RSV replication. Our results correlate with clinical findings where children with RSV had high levels of LDH in nasal washes compared to children with other respiratory viruses (22, 25, 26). Like LDH, caspase increased in the apical compartment during the later phase of infection and correlated significantly with LDH levels (22, 25). However, levels of apoptosis as quantified via caspase did not show significant difference between pediatric and adult HNO-ALIs infected with RSV. This suggests other mechanisms of less regulated cell death such as necrosis can predominate over apoptosis in RSV infection of children compared to adults. In fact, higher levels of apoptosis quantified via caspase 3 and 7 in nasal wash samples have been associated with less severe RSV infection in children (25). Apoptosis is thought to be related to better control of viral replication and less secondary cellular injury as compared to cell death via necrosis or other non-apoptotic pathways (25)

While infectious virus and resultant cell death and damage (caspase and LDH assays) was detected predominantly on the apical compartment, cytokine production was more pronounced in the basolateral compartment. Our HNO-ALIs mimicked the initial cell damage of the airway epithelium caused by local viral effects. Signals from apical RSV infection transmitted through the entire epithelium and eventually into the basolateral compartment, modeling the blood stream. In the infant and adult, this results in the recruitment of the innate and adaptive cellular immune system, which may further augment airway protection and viral control. Overall cytokine release was greater in pediatric HNO-ALIs compared to their adult counterparts, with 5/6 cytokine groups having significantly higher levels compared to adults. Levels of IL-6, IL-8, IL-1α, TNF-α, MIP-1β, MCP-3 and IP10 were significantly higher in pediatric HNOs. Interestingly elevated levels of IL-6, IL-8, and TNF-α in nasopharyngeal samples have been widely correlated with severe RSV disease in infants, including the risk of mechanical ventilation (27, 28). MIP-1β levels in NPA were directly correlated to oxygen therapy (29). However, we and others have demonstrated that high levels of MIP-1β and IP-10 associated with a decreased risk of hospitalization and hypoxia (10, 12). A contrasting finding in adults HNO-ALIs was higher levels of regulatory cytokines. This was driven by IL-17E (IL-25) being significantly higher in the adult HNO-ALIs compared to infants during RSV infection. IL-17E has been associated with inflammatory disorders. More recently, IL-17E is being considered as a barrier cytokine associated with maintaining homeostasis, tissue repair following cellular injury and signaling the immune cells (30). This is contrary to the findings by Petersen et al, where IL17E was linked to increase mucus production and pulmonary inflammation in the RSV mouse model, but it may also have a protective role against viral infections when Th2 cytokines are reduced (31). In general, our findings suggest that pediatric HNO-ALIs produce a more robust cytokine response that may be dysregulated in response to RSV infection.

We also noted 4 biomarkers that were present in higher concentrations in the apical compartment where the apical ciliated cells were infected with RSV and undergoing cellular injury. These were IL-1α, MMP-9, IL-6, and TGF-β. Three of these biomarkers, IL-1α, MMP-9, IL-6, are associated with inflammation and recruitment of inflammatory hemopoetic cells (32–34). In addition, Il-1α functions as an alarmin being released extracellularly during cell necrosis and sequestered during apoptosis. Il-1α is a known trigger of IL-6 and TNF-α from epithelial cells (35). Strikingly, the combined stimulation of Il-6 with TGF-β, both highly released on the apical side of HNO-ALIs, are essential for the generation of IL-17 producing T cells (TH17 cell differentiation) whereas TGF-β stimulation alone is relevant for the differentiation of regulatory T cells (Treg) (36). TH17 cell effector function has both an inflammatory and protective element in mucosal immunity.

Copious mucous production of the upper airway is a hallmark of RSV infection in children. At baseline, pediatric HNO-ALIs appeared to have increased mucus production compared to adult HNO-ALIs. Mucus greatly increased in both adult and pediatric HNO-ALIs during RSV infection. Interestingly, RSV/B/BA infection was associated with greater mucous production compared to RSV/A/ON, while RSV/A/ON caused greater damage to the ciliated apical cells based on LDH release. We are not aware of clinical studies that have evaluated for differences in rhinorrhea between infecting virus subtypes (37) and whether this differences in mucous production and clearance between virus subtypes is clinically relevant. We also found that RSV/A/ON had a more robust overall cytokine response than RSV/B/BA. Overall, we postulate that a balance of cellular damage and appropriately regulated immune response is needed to control the initial upper airway response to RSV infection and prevent viral extension into the lower respiratory tract.

Although our HNO-ALI model has clinical relevance, there are limitations to our findings. Our current HNO-ALI model only examined the upper respiratory tract epithelial response to RSV infection. It will be important to confirm these findings in the human lung organoids representing the lower respiratory tract epithelium of both pediatric and adults. In our study, we compared four adults and four pediatric HNO-ALIs. These numbers are relatively small; however, we were able to detect significant age-specific differences in RSV replication kinetics, cytokine production, and cell damage and recovery between adult and pediatric HNO-ALIs. Currently our HNO-ALI model does not contain immune cells, which could considerably alter how the cells respond to viral load, cytokines, and tissue damage. Future studies with increased complexity of co-culture with immune cells and or endothelium will provide even more physiological results to predict outcomes in children and adults. Although the HNO-ALI model is not a complex model that mimics the pulmonary physiology of animals or humans, it does contain the cellular complexity of the human respiratory epithelium. It is highly permissive to RSV replication unlike most animal models, and the outcome measured such as cytokine response, mucous production, cellular damage in response to RSV infection is similar to the outcomes measured in RSV infected infants (12, 40) and the human challenge model of RSV infection of the upper respiratory tract (38, 39).

In summary, we used the HNO-ALIs to study the initial line of defense against RSV infection. Our data suggests that there are significant differences between the pediatric and adult upper respiratory epithelium in their response to RSV infection with increased inflammatory cytokine response and cell damage in the presence of heightened mucus production. In addition, a significant decrease was observed in the percentage of club cells in pediatric HNO-ALIs compared to adults in response to infection. This can lead to diminished control of inflammation, ciliated cell regeneration, and muco-ciliary clearance with possible extension into the lower respiratory tract. We propose the increased susceptibility to RSV lower respiratory tract infection in infants compared to adults reflects the inability of the infant’s upper respiratory tract epithelium to control RSV infection. Thus, preventive and therapeutic interventions that target RSV infection of the upper respiratory tract may have substantial benefits in reducing lower respiratory tract disease. Lastly, this data could serve as a springboard to understand the increased morbidity and mortality in RSV infection in other at-risk populations not only including young children, but the elderly, patients with asthma, and patients with pulmonary disorders such as chronic obstructive pulmonary disease and cystic fibrosis.

## METHODS

### HNO-ALI cell lines

Eight cell lines of HNO-ALI were established as previously described (9). Briefly, nasal washes and swabs were collected from 4 adults and 4 children under 5 years of age after obtaining signed informed consent by the adult or legal guardian under an approved protocol by Baylor College of Medicine Institutional Review Board. Samples were placed in digestion media [10ml airway organoid (AO) medium + 10mg Collagenase (Sigma C9407) +100μl Amphotericin B], strained to remove debris, and washed several times. After pelleting of the cells and removal of the supernatant, cells were suspended in Matrigel^®^ (Corning, NY) and plated for cell expansion as 3 dimensional (D) HNOs for 3-4 days for pediatric HNOs and approximately 5-7 days for adult HNOs in growth media. HNOs were then enzymatically and mechanically sheared in order to make a single cell suspension, and seeded onto Transwells^®^ (Corning, NY) at a density of 3 × 10^5^ cells/well. HNO-ALIs were maintained in AO media with endothelial growth factor (EGF) (Peprotech-AF-100-15) containing 10μM Y-27632. After 4 days the monolayers of cells were subsequently maintained in an air-liquid environment with differentiation media (PneumaCult-ALI medium from STEMCELL Technologies) added to the lower (basolateral) compartment of the transwells and the epithelium (apical compartment) was air exposed and maintained in a humidified incubator at 37°C with 5% CO2. The differentiation media was replaced every 4 to 5 days. HNO-ALI cultures were maintained for a total of 21 days during which differentiation occured into a pseudostratified multi-cellular ciliated epithelium. At the end of 21 days of differentiation in an air-liquid environment, the HNO-ALI cultures were used for the viral infection studies.

### Study design

Four adult HNO-ALI cell lines (HNO-ALI 2, HNO-ALI 918, HNO-ALI 919, and HNO-ALI 920) and four pediatric HNO-ALI cell lines (HNO-ALI 9003, HNO-ALI 9005, HNO-ALI 9007, and HNO-ALI 9009) were used for this study. The differentiated HNO-ALI cultures were apically infected with RSV/A/USA/BCM813013/2013(ON) (RSV/A/ON), or RSV/B/USA/BCM80171/2010(BA) (RSV/B/BA) at a 0.01 multiplicity of infection (MOI). Two technical replicates of infected HNO-ALI transwells were used for measuring outcomes at each time point: 1, 2, 5, and 8 day post-infection (dpi), except for HNO-ALI 2 which had time points of 1, 2, 5, and 10 dpi. The outcomes measured from the apical and basolateral supernatant samples were virus kinetics, cytokines and chemokines, and biomarkers of cell injury (lactate dehydrogenase) and apoptosis (caspase 3/7), while the membrane-attached epithelium was processed for cell composition and cytopathology.

### Sample collection

Apical wash samples were collected using three consecutive 200 µl washes with AO differentiation media. The combined 600 µl apical wash was diluted 1:1 with 15% glycerol/Iscove media, aliquots were prepared, snapped-frozen and stored at -80°C. All 600 µl of the basolateral media was collected and diluted 1:1 with 15% glycerol/Iscove media, aliquots were prepared, snapped-frozen and stored at -80°C. The 15% glycerol/Iscove media is used to stabilize the virus during freezing and thawing conditions.

### Viral infection

HNO-ALI cell lines were infected with RSV at an MOI of 0.01. Briefly, the virus inoculum (30µl/well) of RSV/A/ON, or RSV/B/BA was added to the apical compartment for 1.5 hours at 36 °C with 5% CO2, and then the inoculum was removed. For mock infection, AO-differentiation media (30 µl/well) alone was added. Similarly, the mock inoculated transwells were incubated for 1.5 hours at 36 °C with 5% CO2, and then the mock inoculum was removed.

### PCR and plaque assays

The viral RNA was extracted using a mini viral RNA kit (Qiagen Sciences) in an automated QIAcube platform according to the manufacturer’s instructions (20). Viral RNA was detected and quantified using real time polymerase chain reaction (RT-PCR) with primers targeting the nucleocapsid gene of RSV/A and RSV/B (20). Infectious RSV (PFU/mL) activity was measured using a quantitative plaque assay as previously described (21). The lower limit of detection was 50 PFU/ml. PFU below the lower limit of detection were assigned a value 1.

### Cytokine and chemokine assay

Cytokines and chemokines secreted by HNO-ALIs in the apical and basolateral compartments were measured and analyzed using the Milliplex cytokine/chemokine magnetic bead panel (Millipore) according to the manufacturer’s instructions. The kits used in this study include (i) the Milliplex human cytokine panel with eotaxin/CCL11, fibroblast growth factor 2 (FGF-2), granulocyte colony stimulating factor (G-CSF), granulocyte-macrophage colony stimulating factor (GM-CSF), interleukin 1 alpha (IL-1α), interleukin 1 beta (IL-1β), interleukin 6 (IL-6), interleukin 8 (IL-8/CXCL8,) interleukin 17E (IL-17E/IL-25), interferon gamma induced protein 10 (IP-10/CXCL10), monocyte chemoattractant protein 1 (MCP-1), monocyte chemoattractant protein 3 (MCP-3), monokine induced by gamma interferon (MIG/CXCL9), macrophage inflammatory protein 1 alpha (MIP1α), macrophage inflammatory protein 1 beta (MIP1β), regulated on activation, normal T cell expressed and secreted (RANTES/CCL5), tumor necrosis factor alpha (TNFα), vascular endothelial growth factor a (VEGF-A), interleukin 33 (IL-33), interferon gamma inducible T-cell alpha chemoattractant (TAC/CXCL11), interleukin 29 or interferon lambda 1 (IL-29/IFN-λ), (ii) the transforming growth factor beta (TGFβ1 Singleplex kit), (iii) the Milliplex human MMP panel 2 with matrix metallopeptidase 9 (MMP9), and matrix metallopeptidase 7 (MMP7), and (iv) the Milliplex human tissue inhibitor of metalloproteinases (TIMP) panel 2 with TIMP1. Data were obtained with Luminex xPONENT for MAGPIX v4.2 build 1324 and analyzed with MILLIPLEX Analyst v5.1.0.0 standard build. All cytokine concentrations less than the lowest standard of each analyte are considered negative, and a value half the concentration of the lowest standard were imputed.

Heat maps were generated showing the expression of each cytokine compared to the mock expression of that cytokine at each time point (1, 2, 5, and 8 dpi). The log2 value of the expression change from mock value of each cytokine in the apical and basolateral compartments was plotted for RSV/A/ON and RSV/B/BA infected HNO-ALIs.

### Lactate dehydrogenase (LDH) and cleaved caspase assays

Total LDH activity was measured in apical wash and basolateral samples using a previously described method (22) . Apical wash and basolateral samples were assayed following protocol instructions (Cytotoxic Detection Kit, Roche Applied Science). To calculate absolute values, L-lactate dehydrogenase (Roche Applied Science) was used to construct a standard curve that demonstrated an ample linear dynamic range (r^2^≥0.998) at the dilutions tested from 0.8 to 110 milliunits per milliliter (mU/mL).

LDH release may be a marker of cell injury or death caused by virus or may be due to apoptosis induced by virus. To measure a marker of apoptosis in apical wash and basolateral samples, the Caspase-Glo-3/7 kit (Promega) was used as previously described (22). The luminescence of caspase was measured with a Synergy H1 microplate reader (Agilent Technologies) and expressed as relative luminescence units (RLU) and converted to units per milliliter (U/mL). A standard curve was constructed that demonstrated an ample linear dynamic range (r2>0.998).

### Immunohistochemistry (IHC) and Immunofluorescence staining

HNO-ALI cell lines were fixed in image-iT™ Fixative Solution (4% formaldehyde) [Catalog number: FB002] for 15 minutes followed by dehydration in ethanol series (30%, 50%, and 70%, each 30 minutes at room temperature or overnight at 4⁰C). The transwell membranes were then embedded in paraffin and sectioned. Standard hematoxylin and eosin (H&E) Periodic acid–Schiff/Alcian-Blue (PAS/AB) staining was performed. For immunofluorescence staining, the sections were deparaffinized in Histo-Clear, followed by washes in an alcohol sequence (100>100>90>70%). Then the slides were rehydrated and exposed to heat-induced antigen retrieval in 10 mM citrate buffer pH (23). The sections were then washed in water for 5 min and blocked for 60 min in 2% bovine serum albumin (BSA) in blocking buffer (PBS). The sections were incubated with the following primary antibodies, keratin 5 (1:2000, KRT5 for basal cells; BioLegend, Catalog number: 905503), SCGB1A1 (1:200, CC10 for club cells; Santa Cruz, Catalog number: sc-9773), acetylated alpha tubulin for cilia (1:1000, Santa Cruz, sc-23950), Mucin 5AC for goblet cells (1:1000, Invitrogen, Catalog number: 45M1), goat polyclonal antibody specific for RSV (1:2000, Abcam, Catalog number: ab20745) overnight at 4 °C. Primary antibodies were washed three times in PBS+0.05% Tween for 10 minutes each, incubated with secondary antibodies (donkey anti-mouse 488 Invitrogen, A-21202, donkey anti-rabbit 568, Invitrogen, A-10042, donkey anti-goat 647, Invitrogen, A-21447) for 2 hours at room temperature, washed twice with PBS, stained with 4′,6-diamidino-2-phenylindole (DAPI), washed twice with PBS, and mounted in Vectashield Plus^®^ (Vector Laboratories, Catalog number H-1900). The slides were stored at −20 °C.

### Immunofluorescence image quantification and analysis

Samples were imaged using high resolution Cytiva DVLive or Olympus IX83 epifluorescence deconvolution microscopes. Images were collected with both a 20x/0.75NA and a 60x/1.42NA objective lenses, with a 10µm z-stack (using optical sections at the recommended Nyquist for each objective). 3D images were deconvolved using a quantitative image restoration algorithm.

Max intensity projections were used for image analysis and processed using Fiji (24). Each experiment had a minimum of three different fields of view quantified as cell counts in Fiji. The average cell count of the different fields were obtained for each specific cell type at the beginning (day 1 and 2), the midpoint (day 5), and the endpoint (day 10 for HNO2, day 8 for all other HNO-ALIs) of infection assays. Counts of goblet cells (Muc5AC+ stained cells), basal cells (Krt5+ cells), club cells (CC10+ cells) after infection with RSV/A/ON and RSV/B/BA were quantified as percentages relative to the total number of DAPI+ cells.

For epithelial area, the epithelium was manually outlined using the draw setting in Fiji and quantified by using the “calculate area” function. The approximate area of apical ciliated cells (acetylated alpha tubulin +) and mucous (Muc5AC+) was quantified using Fiji particle analysis to calculate a total area in pixels. The epithelial area, mucous area, and ciliated area were then converted to μm via using the pixel size information from the imaging specs.

For quantification of IHC images, 5 high-powered field images were taken at 40x/0.65 on a Nikon CiL Brightfield microscope. The percentage of dividing cells (Ki67+ stained cells) were compared to the total number of Hematoxylin+ cells at 1, 2, 5 and 8 dpi.

### Sex as a biological variable

Sex was not considered as a biological variable.

### Statistical Analysis

The study was designed to determine if there were differences between adult and pediatric HNO-ALIs in responses to RSV infection. Study factors were infection condition, time, and cell surface. Outcome variables measured were virus replication, cytokine expression, cell injury, and cell composition. In total, there were 192 dedicated transwells that were used to generate the samples collected under this study design for statistical analysis. These included four adult and 4 pediatric lines each with two technical replicates.

For each outcome variable (except where noted below), a 3×2×4 factorial design approach characterized by infecting condition (mock, RSV/A/ON, RSV/B/BA), cellular compartment (apical or basolateral), and time (1, 2, 5, and 8 dpi) was undertaken to analyze the differences between adult and pediatric HNO-ALIs. Duplicate response values were obtained for most datapoints; therefore, a single observation was represented by the average of the technical replicate response values for analyses. All samples were collected on 1, 2, 5, and 8 dpi except for HNO-ALI 2 that was collected on day 10 and imputed as day 8. Time was treated as a continuous factor to evaluate linear and quadratic effects. Age (adult or pediatric) was included as a binary covariate of primary interest. All two-way interactions with age were included in the model to test whether the age effect differed by infecting condition, cell compartment, and across time. Highly non-significant interaction terms were dropped one at a time from the model to re-evaluate the model with remaining interaction terms. A 3-way interaction term (age-virus-time) was only introduced in the model if it was shown that there was some indication of significance (liberal threshold of p<0.20) in at least two of the three 2-way interactions. For a significant interaction term, the emmeans package (1.8.7) with the contrast function was used to generate contrast tests for comparing the difference in levels of one factor (age effect) with each of the levels in the other factor. No multiple comparison adjustment procedure was applied. Point estimates (beta coefficients, odds ratios, relative risks) were reported along with corresponding 95% confidence interval (CI) and statistical significance was assigned when p values were <0.05. Statistical analyses were performed with R software v4.3.2 (R Foundation for Statistical Computing).

#### Virus replication

Ordinary least squares linear regression was used to analyze the age effect on viral kinetics of RSV/A/ON and RSV/B/BA as measured by PCR log10 copy numbers from the apical wash samples and by plaque assay log10 PFU values from apical wash samples, over 4 timepoints.

#### Cytokine expression

We evaluated 24 different cytokines in both the apical and basolateral compartments from 8 different HNO-ALIs infected with RSV/A/ON, RSV/B/BA or mock over 4 timepoints. To account for the differing magnitudes and variances of the cytokine responses, the raw cytokine concentration values were log transformed and then standardized by converting to Z-scores. For each cytokine, a Z-score was calculated by subtracting the overall average concentration from the individual concentration value and dividing that result by the standard deviation of all concentration values within that single cytokine. A composite Z-score was calculated for each cytokine functional group by taking the average of the Z-scores of the respective cytokines within a cytokine group. These composite Z-scores were then converted into probabilities from a standard normal cumulative distribution function using the pnorm function in R. A separate fractional logit regression was performed on each cytokine functional group’s composite Z-score probabilities based on quasi-binomial generalized linear model. Fractional logit regression implements quasi-maximum likelihood estimators with robust standard errors fitting a model conditional on the set of independent variables (infection condition, cell compartment, time, age and 2-way interactions) providing estimates in odds ratios (OR).

#### Cell injury

LDH and caspase concentration values were log transformed, standardized to Z-scores, converted to probabilities, and were analyzed with fractional logit regression. LDH and caspase concentration values were not available for HNO-ALI 2, and caspase concentration values were not available for HNO-ALI 9009 for RSV/A/ON infection.

#### Cell composition

Lastly, we quantitated the percent of basal cells, goblet cells, secretory cells, and proliferative cells as well as cilia, mucous and epithelial area in the HNO-ALIs over time describing cell type specific changes during RSV infection. Percentages of basal, goblet, secretory, and proliferative cells were rescaled to the range 0-1 by dividing by 100 for use in fractional logit regression. Poisson regression analysis was performed to analyze cilia, mucous and epithelial area to obtain robust standard errors for the risk ratio (RR) estimates.

### Study Approval

Nasal washes and swabs to generate nose organoids were collected from adults and children after obtaining signed informed consent from the adult or legal guardian under an approved protocol by the Institutional Review Board at Baylor College of Medicine.

## Data Availability

A Supporting Data Values file with all reported data values will be available as part of the supplemental material.

## Author contributions

GMA designed the project, performed experiments, analyzed data, and wrote the manuscript draft. DN, AMW, EMS, AR, LA, and TM performed infections, viral kinetics and cytokine measurements, and imaging, and edited the manuscript. EN consented subjects, obtained nasal washes from subjects, and edited the manuscript. DH, LFS performed the statistical analysis and edited the manuscript. AK provided the organoid cultures and edited the manuscript. HLJ, EM, and FS performed microscopy and edited the manuscript. SEB supervised established organoid cultures and edited the manuscript. PAP and VA designed the project, analyzed data, obtained funding, and wrote and edited the manuscript. All authors reviewed and approved the final manuscript.

## ACKNOWLEDGMENTS

This work was supported by funds from the National Institutes of Health (NIH) grants U19 AI116497 (SB, VA, PAP) and U19AI144297 (SB, VA, PAP). Imaging for this project was supported by the Integrated Microscopy Core at Baylor College of Medicine and the Center for Advanced Microscopy and Image Informatics (CAMII) with funding from NIH (DK56338, CA125123, ES030285, S10OD030414), and CPRIT (RP150578, RP170719)

## Conflict of interest

The authors declare no conflicts of interest.

---

**Figure S1.**
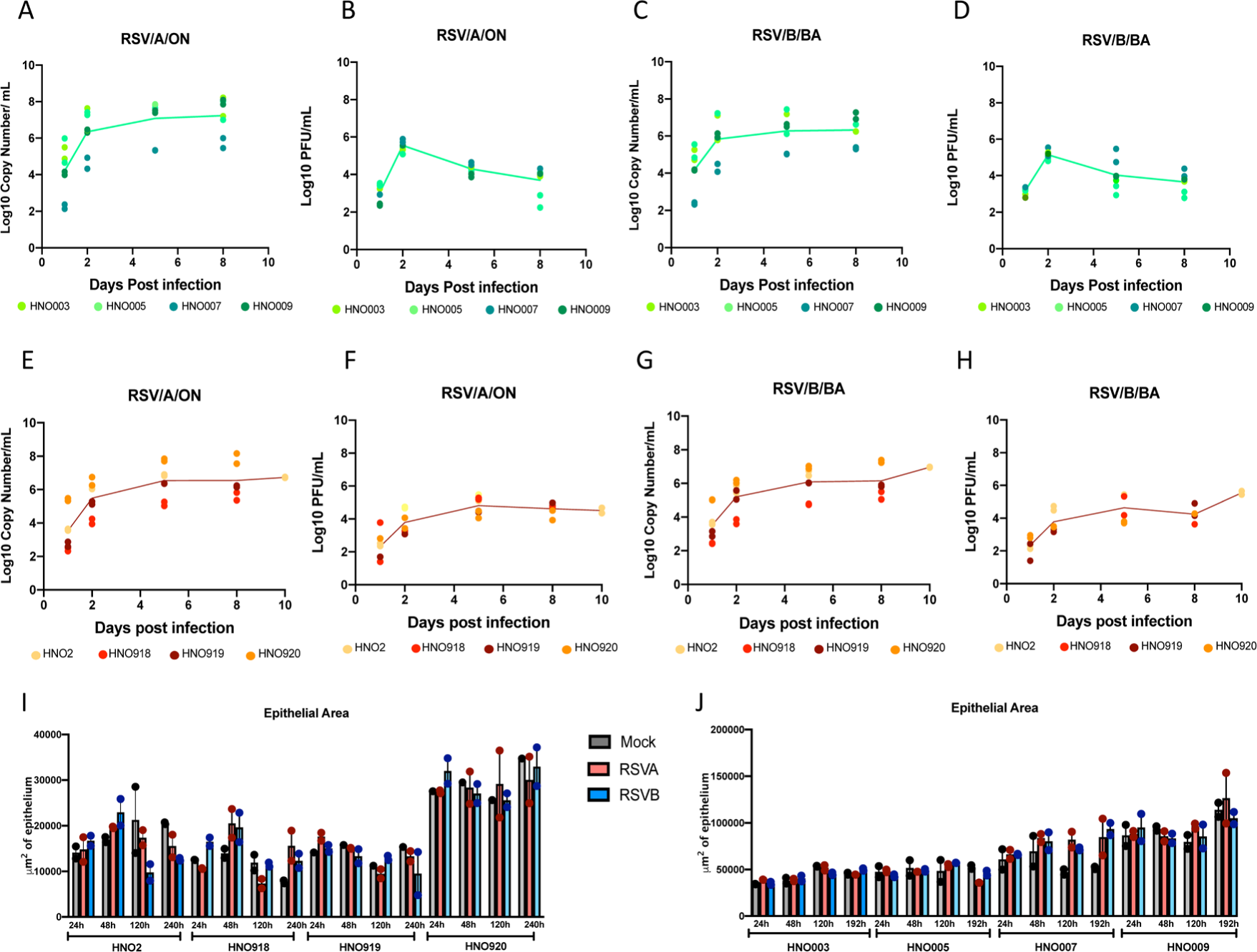
Replication kinetics and morphologic analysis of RSV infected individual pediatric HNO-ALIs and adult HNO-ALIs. **A)** RT-PCR for apical RSV/A/ON for 4 pediatric HNO-ALIs at 1, 2, 5, and 8 days post-infection (dpi). Each HNO-ALI is assigned a specific color, and two replicates of each line are shown on the graph, with composite mean values plotted in a line. **B)** Corresponding plaque data for RSV/A/ON pediatric HNO-ALIs. **C)** RT PCR for apical RSV/B/BA for 4 pediatric HNOs at 1, 2, 5, and 8 dpi. **D)** Corresponding plaque data for RSV/B/BA pediatric HNO-ALIs. **E)** RT-PCR for apical RSV/A/ON for 4 adult HNO-ALIs at 1, 2, 5, and 8 or dpi. Each HNO-ALI is assigned a specific color, and two replicates of each line are shown on the graph, with composite mean values plotted in a line. **F**) Corresponding plaque data for RSV/A/ON adult HNO-ALIs. **G)** RT PCR for apical RSV/B/BA for 4 adult HNO-ALIs at 1, 2, 5, and 8 dpi. **H)** Corresponding plaque data for RSV/B/BA adult HNO-ALIs. **I)** Area of the epithelium of 4 adult HNO-ALIs after infection with RSV/A/ON, RSV/B/BA, or mock infection at 1, 2, 5, and 8 dpi. **J)** Area of the epithelium of 4 pediatric HNO-ALIs after infection with RSV/A/ON, RSV/B/BA, or mock infection at 1, 2, 5, and 8 dpi.

**Figure S2.**
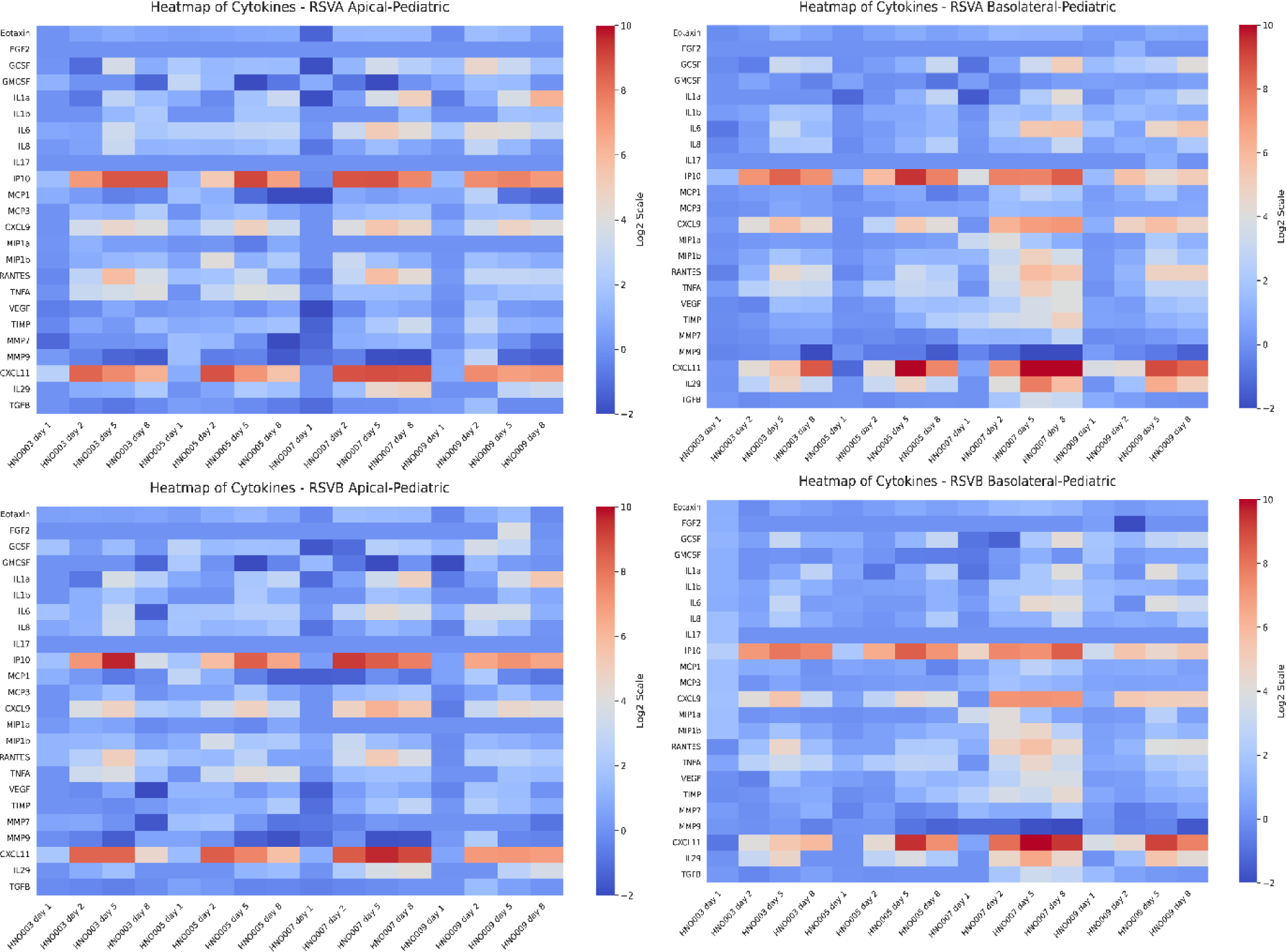
Heat map of cytokine expression for all 4 pediatric HNO-ALI lines. Expression over time for RSV/A/ON apical, RSV/A/ON basolateral, RSV/B/BA apical, and RSV/B/BA basolateral. Each value is normalized to the corresponding mock value at that time point.

**Figure S3.**
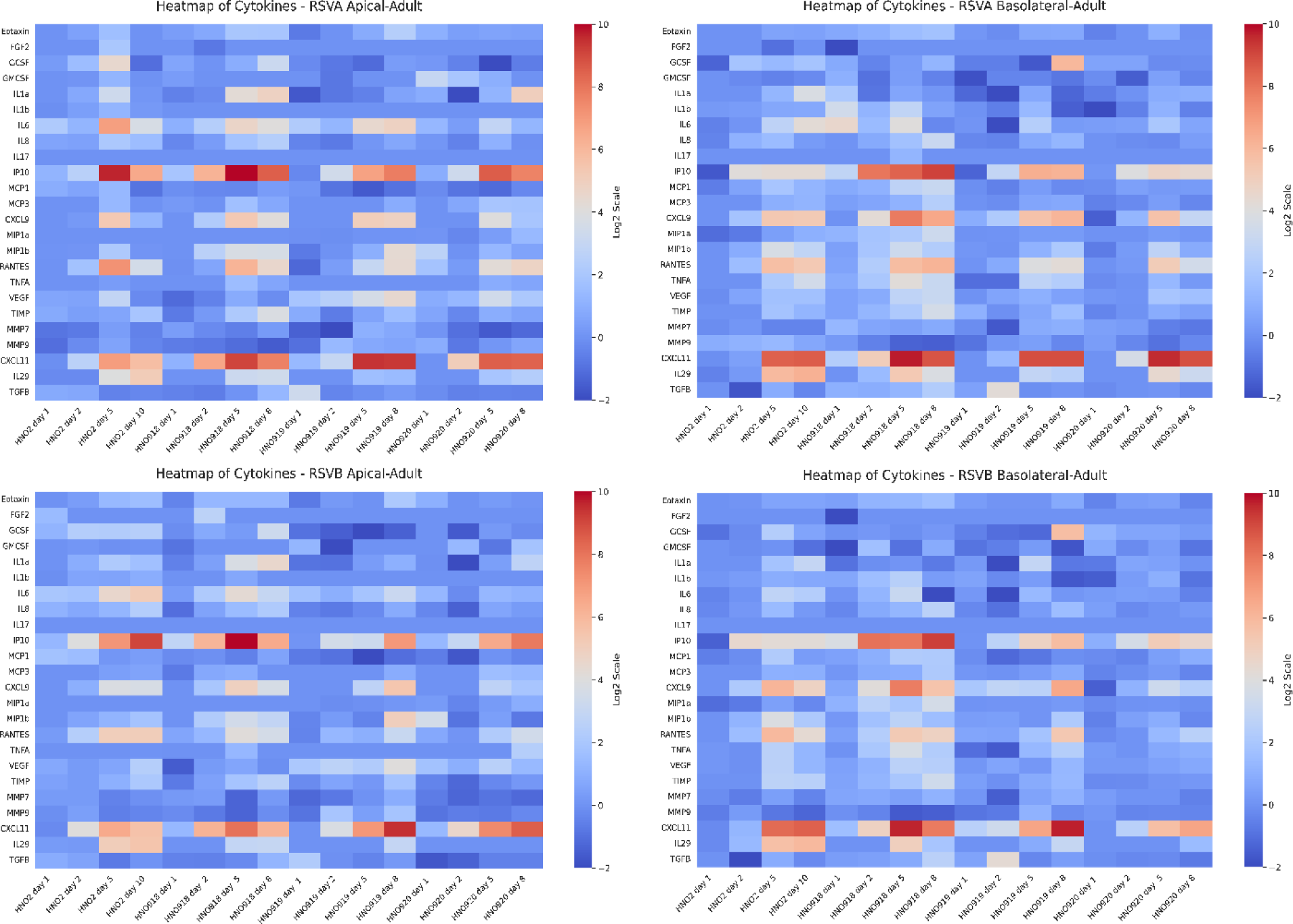
Heat map of cytokine expression for all 4 adult HNO-ALI lines. Expression over time for RSV/A/ON apical, RSV/A/ON basolateral, RSV/B/BA apical, and RSV/B/BA basolateral. Each value is normalized to the corresponding mock value at that time point.

**Figure S4.**
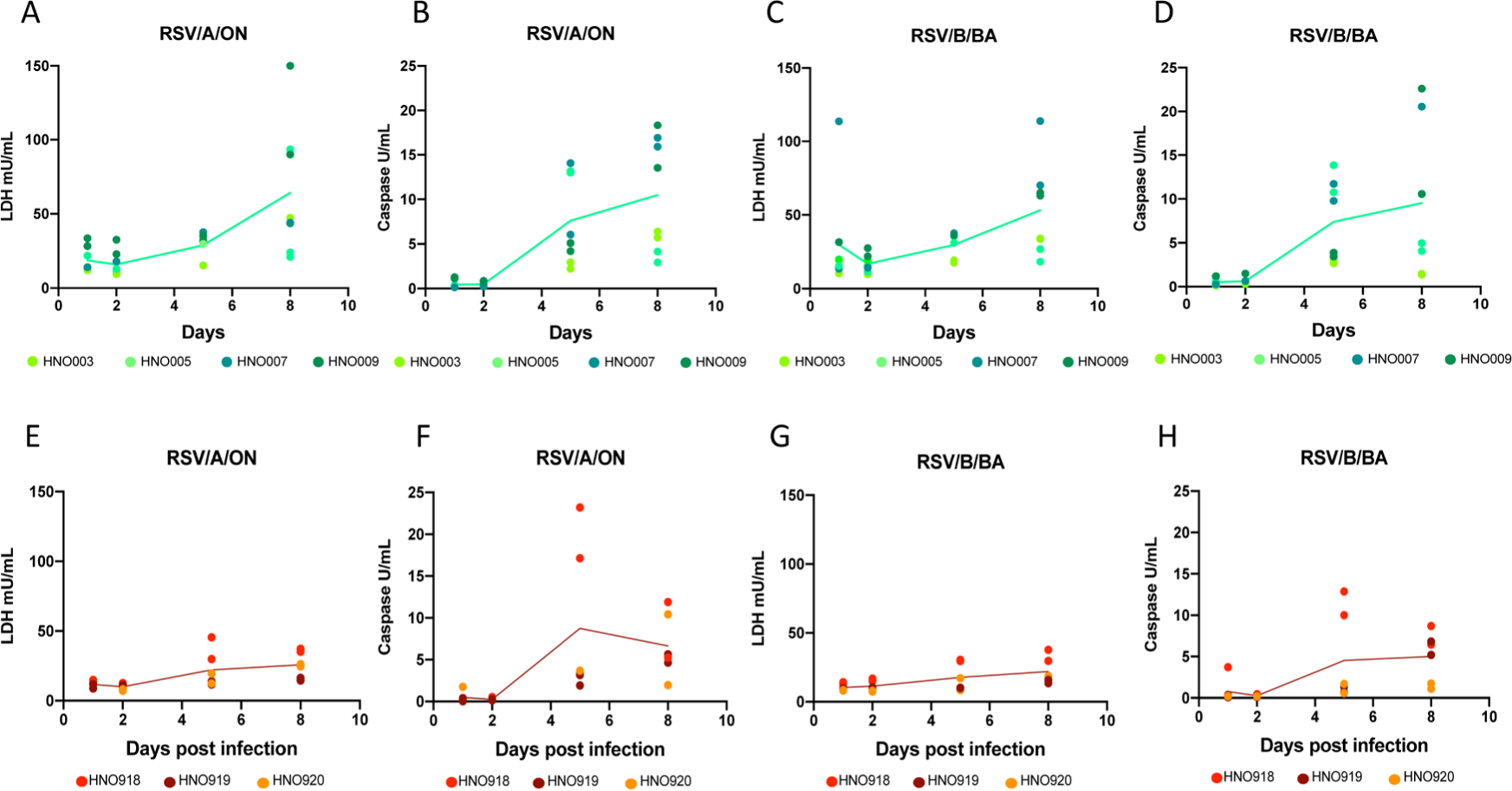
Apical LDH and Caspase levels during RSV infection in individual pediatric and adult HNO-ALIs. **A)** Levels of apical LDH for 4 pediatric HNO-ALIs at 1, 2, 5, and 8 days post-infection (dpi) with RSV/A/ON. Each HNO-ALI is assigned a specific color, and two replicates of each line are shown on the graph, with composite mean values plotted in a line. **B)** Levels of apical caspase for 4 pediatric HNO-ALIs at 1, 2, 5, and 8 dpi with RSV/A/ON. **C)** Levels of apical LDH and **D)** Levels of apical caspase for 4 pediatric HNO-ALIs at 1, 2, 5, and 8 dpi with RSV/B/BA. **E)** Levels of apical LDH and **F)** and levels of apical caspase for 3 adult HNO-ALIs at 1, 2, 5, and 8 dpi with RSV/A/ON. **G)** Levels of apical LDH and **H)** level of apical caspase for 3 adult HNO-ALIs at 1, 2, 5, and 8 dpi with RSV/B/BA.

**Figure S5.**
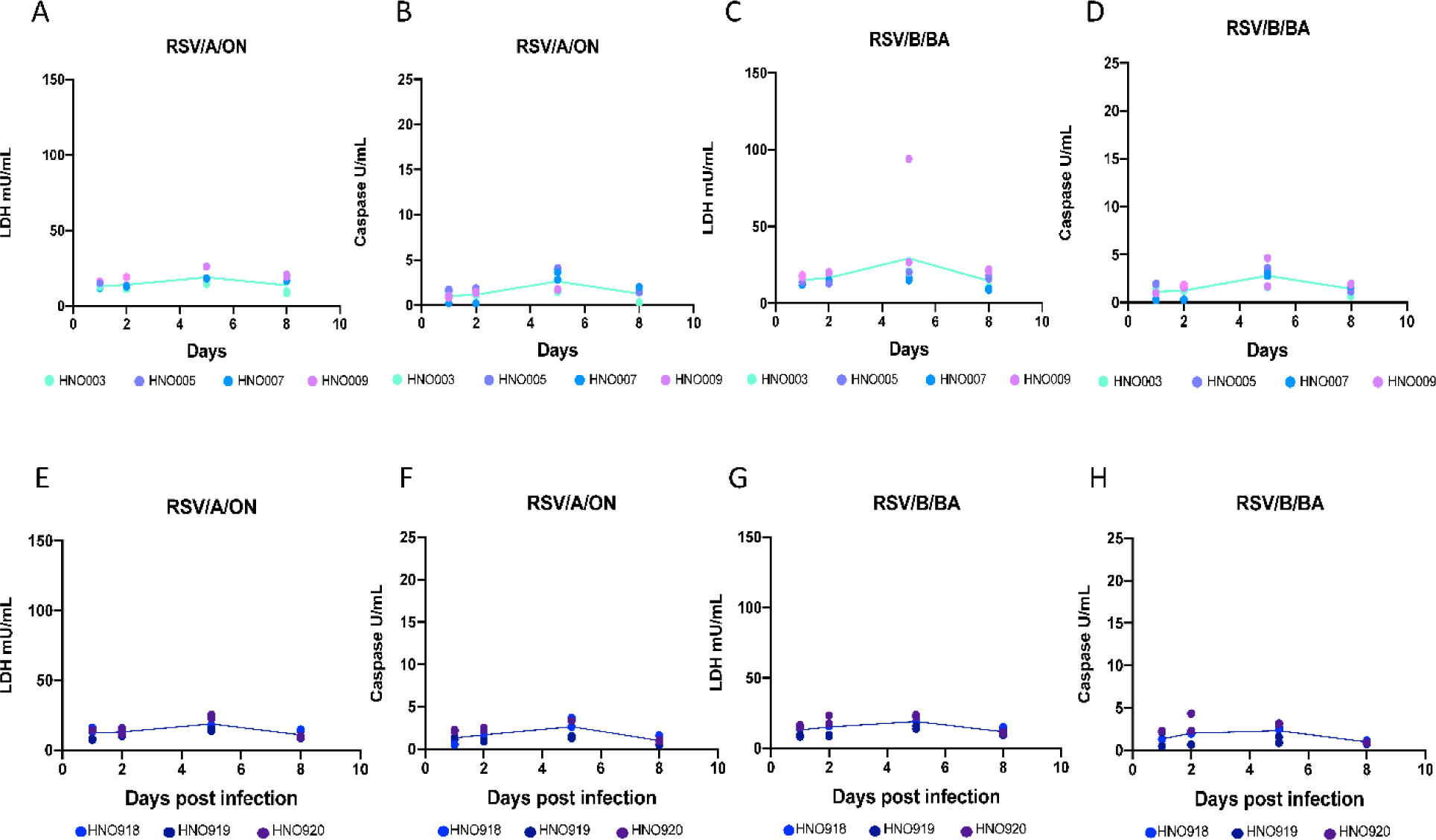
Basolateral LDH and Caspase levels during RSV infection in individual pediatric and adult HNO-ALIs. **A)** Levels of basolateral LDH for 4 pediatric HNO-ALIs at 1, 2, 5, and 8 days post-infection (dpi) with RSV/A/ON. Each HNO-ALI is assigned a specific color, and two replicates of each line are shown on the graph, with composite mean values plotted in a line. **B)** Levels of basolateral caspase for 4 pediatric HNO-ALIs at 1, 2, 5, and 8 dpi with RSV/A/ON. **C)** Levels of basolateral LDH and **D)** Levels of basolateral caspase for 4 pediatric HNO-ALIs at 1, 2, 5, and 8 dpi with RSV/B/BA. **E)** Levels of basolateral LDH and **F)** and levels of basolateral caspase for 3 adult HNO-ALIs at 1, 2, 5, and 8 dpi with RSV/A/ON. **G)** Levels of basolateral LDH and **H)** level of basolateral caspase for 3 adult HNO-ALIs at 1, 2, 5, and 8 dpi with RSV/B/BA.

**Figure S6.**
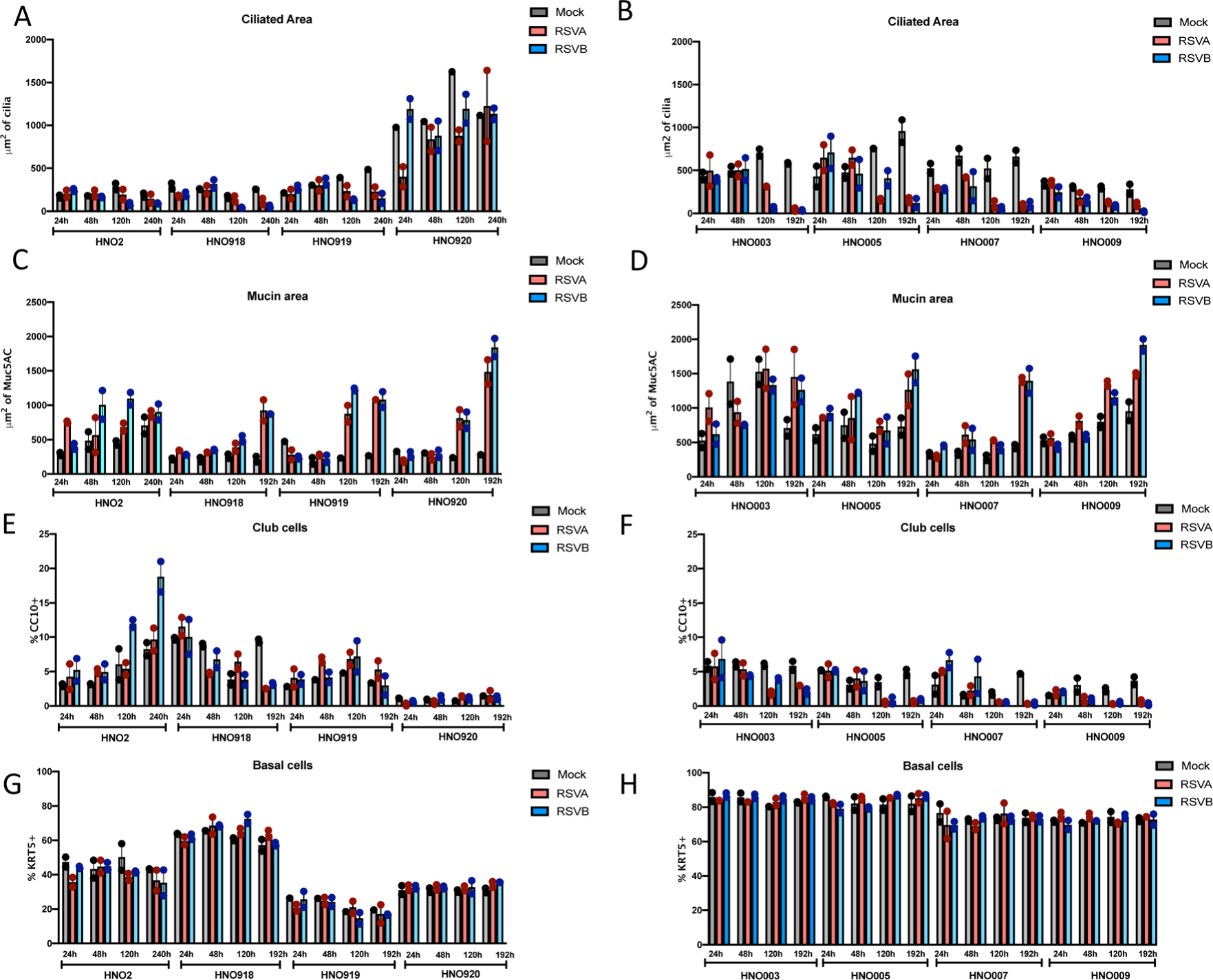
Cellular and morphological changes in pediatric and adult HNO-ALIs. **A)** Amount of apical ciliated area after infection with RSV in 4 adult HNO-ALIs and **B)** 4 pediatric HNO-ALIs at 1, 2, 5, and 8 days post-infection (dpi). with RSV/A/ON, RSV/B/BA, or mock infection. **C)** Area of mucous quantified by Muc5AC+ after infection with RSV in 4 adult HNO-ALIs and **D)** 4 pediatric HNO-ALIs at 1, 2, 5, and 8 dpi. **E)** Percentage of CC10+ club cells over / number of DAPI+ cells after infection with RSV in 4 adult HNO-ALIs and **F)** 4 pediatric HNO-ALIs at 1, 2, 5, and 8 dpi. **G)** Percentage of Krt5+ basal cells /total number of DAPI+ cells after infection with RSV in 4 adult HNO-ALIs and **H)** and 4 pediatric HNO-ALIs at 1, 2, 5, and 8 dpi.

## Notes

### Competing Interest Statement

The authors have declared no competing interest.

